# The proliferative history shapes the DNA methylome of B-cell tumors and predicts clinical outcome

**DOI:** 10.1101/2020.02.06.937383

**Authors:** Martí Duran-Ferrer, Guillem Clot, Ferran Nadeu, Renée Beekman, Tycho Baumann, Jessica Nordlund, Yanara Marincevic-Zuniga, Gudmar Lönnerholm, Alfredo Rivas-Delgado, Silvia Martin, Raquel Ordoñez, Giancarlo Castellano, Marta Kulis, Ana Queirós, Lee Seung-Tae, Joseph Wiemels, Romina Royo, Montserrat Puiggrós, Junyan Lu, Eva Gine, Sílvia Beà, Pedro Jares, Xabier Agirre, Felipe Prosper, Carlos López-Otín, Xosé S. Puente, Christopher C. Oakes, Thorsten Zenz, Julio Delgado, Armando López-Guillermo, Elías Campo, José Ignacio Martin-Subero

## Abstract

We report a systematic analysis of the biological and clinical implications of DNA methylation variability in five categories of B-cell tumors derived from B cells spanning the entire maturation spectrum. We used 2056 primary samples including training and validation series and show that 88% of the human DNA methylome is dynamically modulated under normal and neoplastic conditions. B-cell tumors display both epigenetic imprints of their cellular origin and *de novo*, disease-specific epigenetic alterations that in part are related to differential transcription factor binding. These differential methylation patterns were used by a machine-learning approach to create a diagnostic algorithm that accurately classifies 14 B-cell tumor entities and subtypes with different clinical management. Beyond this, we identified extensive patient-specific epigenetic variability targeting constitutively silenced chromatin regions, a phenomenon we could relate to the proliferative history of normal and neoplastic B cells. We observed that, depending on the maturation stage of the tumor cell of origin, mitotic activity leaves different imprints into the DNA methylome. Subsequently, we constructed a novel DNA methylation-based mitotic clock called epiCMIT (<u>epi</u>genetically-determined <u>C</u>umulative <u>MIT</u>oses), whose lapse magnitude represents a strong independent prognostic variable within specific B-cell tumor subtypes and is associated with particular driver genetic alterations. Our findings reveal DNA methylation as a holistic tracker of B-cell tumor developmental history, with implications in the differential diagnosis and prediction of the outcome of the patients.

## Introduction

The process of neoplastic transformation implies a dramatic alteration of cellular identity^1^. However, cancer cells partially maintain molecular imprints of the cellular lineage and maturation stage from which they originate^2^. B-cell neoplasms are a paradigmatic model of this phenomenon, as the maturation stage of different B-cell neoplasms is the main principle behind the World Health Organization classification of these tumors^3^. Recent studies have focused on the analysis of the DNA methylome, a *bona fide* epigenetic mark related to cellular identity and gene regulation^4,5^ during the entire B-cell maturation program^6^ and in various B-cell neoplasms spanning the entire maturation spectrum. These include B-cell acute lymphoblastic leukemia (ALL)^7,8^ derived from precursor B cells, mantle cell lymphoma (MCL)^9^ and chronic lymphocytic leukemia^10,11^(CLL) derived from pre- and post-germinal center mature B cells, diffuse large B-cell lymphoma (DLBCL)^12,13^ derived from germinal center B cells, and multiple myeloma (MM)^14,15^ derived from terminally-differentiated plasma cells. These studies have revealed a dynamic DNA methylome during B-cell maturation as well as novel insights into the cellular origin, pathogenic mechanisms and clinical behavior of B-cell neoplasms reviewed in^16^. However, a global analysis of the entire normal cell differentiation program and derived neoplasms is neither available for B cells nor for any other human cell lineage. Thus, we herein exploit both previously generated DNA methylation datasets as well newly generated data to systematically decipher the sources of DNA methylation variability across B-cell neoplasms. This comprehensive approach using over 2000 samples reveals previously hidden biological insights and clinical associations. In particular, *de novo* disease-specific hypomethylation in active regulatory regions is associated with differential transcription factor binding and targets genes important for disease-specific pathogenesis. From the clinical perspective, we define a set of epigenetic biomarkers that can accurately classify B-cell neoplasms requiring differential clinical management and construct a DNA methylation-based mitotic clock called epiCMIT as a personalized predictor of clinical behavior within each B-cell neoplasm.

## Results

### Shared DNA methylation dynamics in normal and neoplastic B cells

We analyzed previously published DNA methylation profiles of samples from normal and neoplastic B cells spanning the entire B-cell differentiation spectrum, all generated with the 450k microarray platform from Illumina. These included 10 normal B cell subpopulations^6^ as well as the five common categories of B-cell neoplasms, i.e. ALL^7,8^, MCL^9^, CLL^10,17^, DLBCL (own unpublished series) and MM^14^ (Fig. 1a and Supplementary Table 1). Following the guidelines of the TCGA Consortium, we selected samples containing a tumor-cell content more than 60% https://www.cancer.gov/about-nci/organization/ccg/blog/2018/bcr-tips. This proportion of tumor cells was estimated by flow cytometry^6,9,10,14,17^, genetic data^18^ and/or lineage-specific DNA methylation patterns (Supplementary Table 2). To validate that 60% was an appropriate threshold for B-cell tumors, we analyzed 5 MCLs and 3 CLLs in which we had DNA extracted both from sorted (minimum purity of 86%) and unsorted tumor cells (purity between 48% and 77%). Unsupervised analyses showed that samples tightly clustered according to patient number and not based on their tumor cell content (Extended Data Fig. 1a). In fact, tumor cell content in these paired samples was reflected in a minor component accounting for only 2% of the total variability, and therefore was considered negligible (data not shown). Tumor cell content estimated by any of the three methods was in general highly correlated (Extended Data Fig. 1b) and in most of the cases, we used DNA methylation data to estimate tumor purity. However, MM samples showed that DNA methylation-based estimation of tumor cell content was far lower than that estimated by flow cytometry (Extended Data Fig. 1c). This result is congruent with previous DNA methylation data indicating that MM losses the B cell identity^14^. Unexpectedly, some DLBCL samples also showed a similar effect (Extended Fig. 1c), and therefore in MM and DLBCL, tumor cell content was estimated by flow cytometry and genetic data, respectively. After all filtering criteria (Methods), we generated a curated data matrix containing 1595 high quality samples (Fig. 1a and Supplementary Table 1) with DNA methylation values for 437182 CpGs, which was used in all downstream analyses.

**Fig. 1:**
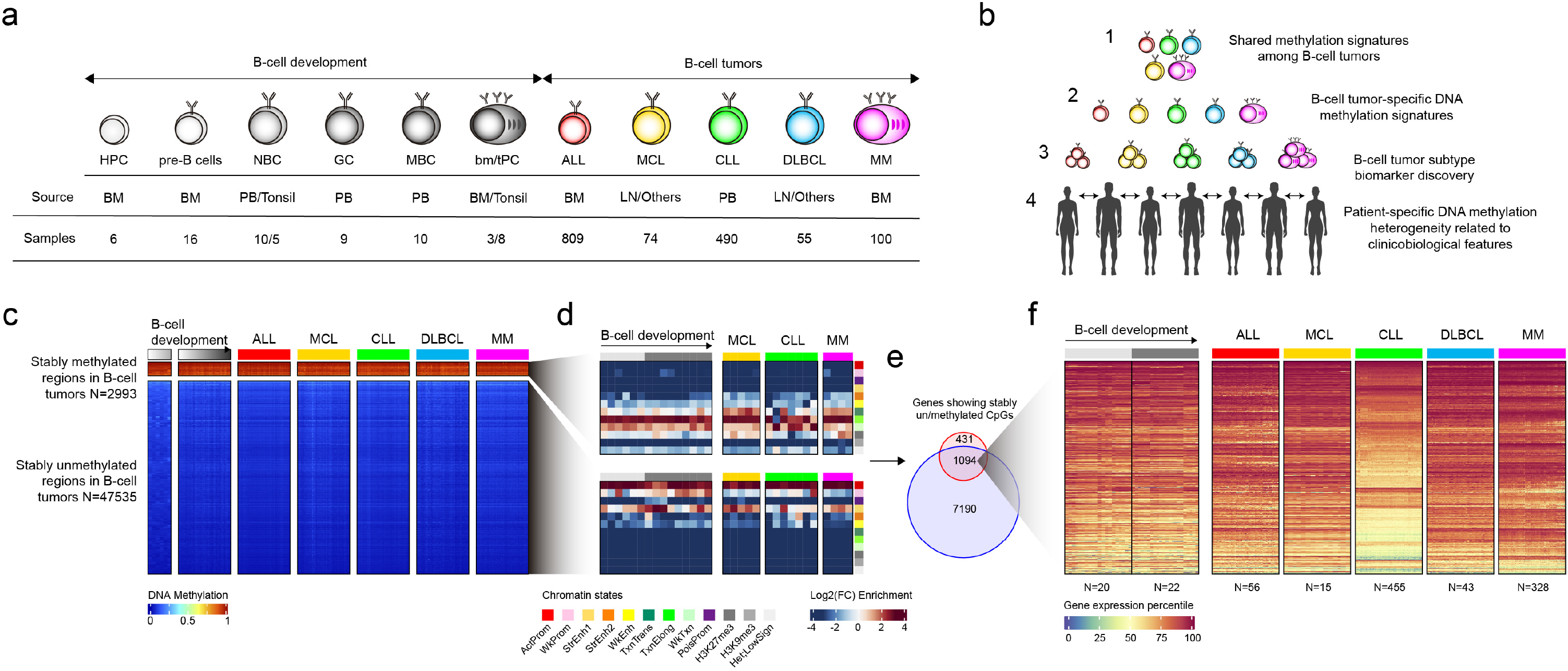
Shared DNA methylation dynamics in B-cell tumors. **a,** Number of normal and neoplastic B-cells included in the study with tumor cell content of at least 60% (additional samples for validations are used in Fig. 3 and Fig. 6, which are later detailed). HPC, hematopoietic precursor cells; pre-B, precursor B-cell and immature B cells; NBC, naïve B cells; GC, germinal center B cells; MBC, memory B cells; tPC, tonsillar plasma cells; bmPC, bone-marrow plasma cells; ALL, acute lymphoblastic leukemia; MCL, mantle cell lymphoma; CLL, chronic lymphocytic leukemia; DLBCL, Diffuse large B cell lymphoma; MM, multiple myeloma; BM, bone marrow; PB, peripheral blood; LN, lymph node. **b**, Different levels of DNA methylation variability addressed in the study. **c,** Heatmaps showing shared DNA hyper- (top) and hypomethylation (bootm) in B-cell tumors. **d,** Chromatin state enrichments of regions sharing DNA hyper- and hypomethylation (all CpGs of 450k array were used as background). ActProm, Active promoter; WkProm, Weak promoter; StrEnh1, Strong enhancer 1 (promoter-related); StrEnh2, Strong enhancer 2; WkEnh, Weak enhancer; TxnTrans, Transcription transition; TxnElong, Transcription elongation; WkTxn, Weak transcription; PoisProm, Poised promoter; H3K27me3, Polycomb-repressed region; H3K9me3, H3K9me3 heterochromatin; Het;LowSign, Het;LowSign heterochtomatin. **e,** Overlap of the target genes of stably methylated and unmethylated CpGs. **f,** Gene expression percentile within each sample of genes showing stable hyper- and hypomethylation.

This large comprehensive dataset offered an exceptional opportunity to stepwise dissect the DNA methylation variability of normal and neoplastic B cells at different magnitude levels, including cancer-specific, tumor entity-specific, tumor subentity-specific and individual-specific variability (Fig. 1b). Before studying such DNA methylation dynamics, we identified that only 12% of the studied CpGs show stable DNA methylation levels in normal and neoplastic B cells (Fig. 1C and Supplementary Table 3). We characterized these CpGs based on genetic location, CpG content and chromatin states from primary samples from the Blueprint consortium^19^. We identified that stably methylated regions were located at gene-bodies of actively transcribed genes whereas stably unmethylated sites were prevalent in CpG islands of promoter regions showing active chromatin marks (Fig. 1d, e). Stably methylated and unmethylated CpGs mainly converged into the same genes (Fig. 1e and Extended Data Fig. 1e), which are highly expressed in normal and neoplastic B cells (Fig. 1f) and are involved in cellular functions such as cell cycle, RNA processing and energy metabolism (Extended Data Fig. 1g). These results indicate, on the one hand, that the stable fraction of the DNA methylome affects genes involved in fundamental cellular functions showing an epigenetic signature of expressed genes (Extended Data Fig. 1g). On the other hand, they show that the great majority of the DNA methylome (88%) is labile during normal B cell differentiation and neoplastic transformation.

We next wondered whether all B-cell tumors, regardless the entity, share a unifying DNA methylation signature related to their neoplastic nature. As 25% of the DNA methylome is modulated during normal B cell differentiation^6,11^, we took the remaining stable 75% and compared the methylomes of normal and neoplastic B cells. This analysis did not identify any consistent *de novo* DNA methylation signature that is shared by all B-cell neoplasms under study. Instead, DNA methylation variability is related to differences among B-cell tumor entities and subtypes as well as patientspecific variability, as will be shown in the following sections.

### Disease-specific hypomethylation targeting regulatory regions is associated with specific transcription factor bindings and differential gene expression

We next aimed at studying whether disease-specific DNA methylation patterns may unravel differential pathogenic mechanisms underlying each B-cell neoplasm. Overall, a principal component analysis (PCA) showed that different B-cell neoplasms form distinct clusters (Fig. 2a). A further analysis indicated that the first nine components contained information regarding specific B-cell neoplasms and allowed us to distil the main biological sources of DNA methylation variability (Extended Data Fig 2a). The first component was related to B-cell development, separating neoplasms according to the maturation stage of their cellular origin, i.e. ALL together with pre-germinal center B cells and mature B cell neoplasms together with germinal center B cells, memory B cells and plasma cells. The remaining components showed tumor-specific patterns, such as PC3, PC7 and PC9 related to MM, DLBCL and MCL-specific variability, respectively. Next, in order to identify DNA methylation signatures specifically associated with malignant transformation, we focused our next analysis on the genome fraction with stable (i.e. B-cell independent) DNA methylation levels across B-cell differentiation^6^ (Fig. 2b). We showed varying numbers of tumor-specific DNA methylation (tsDNAm) changes, ranging from 616 in CLL to 49279 in MM (Fig. 2b, Supplementary Tables 4 and 5). Remarkably, we observed that DNA methylation changes manifested differently in distinct neoplasms. Overall, hypermethylation was enriched at CpG islands and promoter related regions, whereas hypomethylation at low CpG content regions such as open sea, shore and shelfs (Extended Data Fig. 2c). ALL and DLBCL showed more tumor-specific DNA hypermethylation (tsDNAm-hyper) whereas MCL, CLL and MM acquired more tumor-specific DNA hypomethylation (tsDNAm-hypo), being this skew towards hypomethylation remarkable in MM (Fig. 2b-c and Extended Data Fig. 2b). To shed light into the potential causes of this phenomenon, we analyzed the expression levels of DNA methyltransferases (DNMTs) during normal B-cell maturation^6^. We identified that DNMT1, which has been linked both to DNA methylation maintenance and DNA hypermethylation of CpG islands marked by polycomb^20^, shows the highest expression levels in precursor B cells and germinal center B cells, the respective cells of origin of the mostly hypermethylated ALL and DLBCL. Conversely, bone marrow plasma cells, the cell of origin of the mostly hypomethylated MM, showed the lowest levels of DNMT1 expression (Extended Data Fig. 2d). Although the precise mechanisms remain to be elucidated, the level of expression of DNMTs of the respective cells of origin may influence how DNA methylation changes are manifested in distinct B-cell neoplasms.

**Fig. 2:**
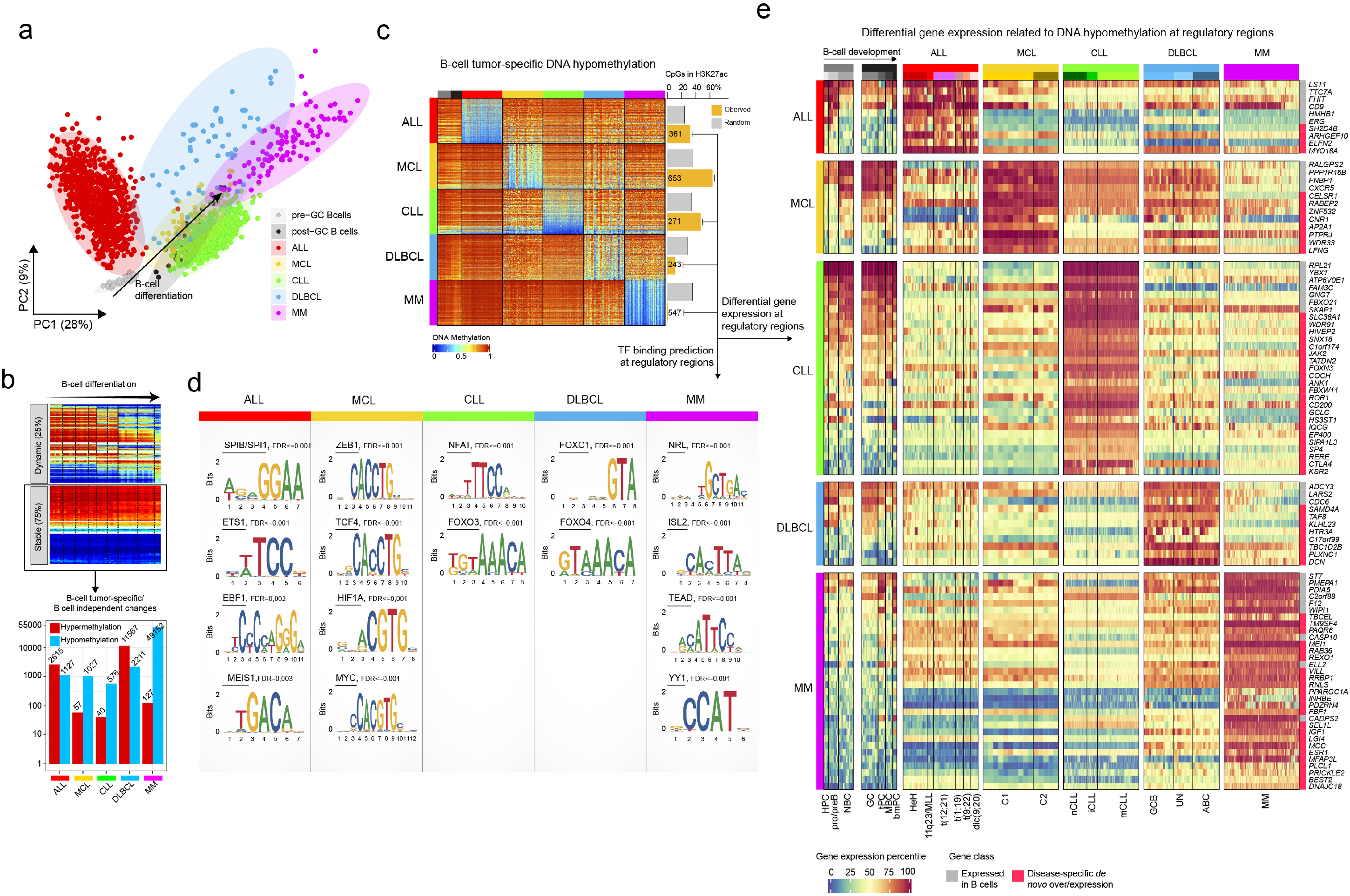
Disease-specific hypomethylation at H3K27ac regions is associated with specific transcription factor bindings and differential gene expression. **a,** Principal component analysis of normal and neoplastic B-cells, including ALL, MCL, CLL, DLBCL and MM. ALL, acute lymphoblastic leukemia; MCL, mantle cell lymphoma; CLL, chronic lymphocytic leukemia; DLBCL, Diffuse large B cell lymphoma; MM, multiple myeloma. **b,** Number of *de novo* DNA methylation changes in each B-cell tumor entity. **c,** Heatmap showing entity-specific hypomethylation and the number of CpGs falling at regulatory regions. Per each B-cell tumor, the same number of *de novo* CpGs was randomly chosen from the 450K array and interrogated the percentage falling at regulatory regions. **d,** Transcription factor binding site predictions for *de novo* hypomethylated CpGs falling at regulatory regions in c, for each B-cell tumor. **e,** Differential gene expression percentiles among B-cell tumors for genes showing specific-hypomethylation at regulatory regions.

Transcription factors (TFs) have been reported to affect DNA methylation levels of regulatory regions upon their binding to the DNA^21,22^. Therefore, we performed TFs binding site prediction analysis in active regulatory elements (i.e. marked by H3K27ac) containing tsDNAm-hypo CpG (Fig. 2d and Methods). Interestingly, the entities in which tsDNAm-hypo was predominantly located in H3K27ac regions (Fig. 2c) showed enrichments for binding sites of TFs expressed in each respective entity (Extended Data Fig. 2e and Supplementary Table 6) and with a previously reported implication in their pathogenesis, such as SPI1/SPIB and EBF1 in ALL, TCF/ZEB in MCL, and NFAT in CLL (Fig. 2d)^23–25^. In the case of DLBCL and MM, their associated tsDNAm-hypo CpGs were actually depleted of regulatory elements (Fig. 2c), suggesting that TF binding may not be a major factor leading to their tumor-specific DNA methylation signatures. However, focusing only on the underrepresented H3K27ac-containing tsDNAm-hypo CpGs, we could also detect significant relationships with TFs potentially involved in the respective diseases, such as FOX family in DLBCL^26^, and NRL (a member of the oncogenic MAF family), ISL1, TEAD, and YY1 in MM^27–30^.

Beyond the potential role of TFs in shaping tumor-specific DNA methylation signatures, we also investigated the downstream transcriptional consequences of tsDNAm-hypo signatures. An analysis of transcriptional profiles of cases from all five diseases revealed a total of 94 genes associated with tsDNAm-hypo genes expressed in a disease-specific manner (Fig. 2e). Although some of the identified genes have been shown to be specifically expressed in a particular disease, such as *CTLA4* and *KSR2* in CLL^31^, this comprehensive analysis provides a rich resource of novel epigenetically-regulated candidate oncogenes for mechanistic studies in each B-cell neoplasm entity.

### Accurate classification of 14 clinico-biological subtypes of B cell neoplasms using epigenetic biomarkers

The B-cell neoplasms shown in Fig. 1a represent broad categories which are further classified into subtypes with different clinico-biological features based on genetic, transcriptional or epigenetic features^3^. These include high-hyperdiploid (HeH), 11q23/MLL, t(12;21), t(1;19), t(9;22) and dic(9;20) ALLs^7^; C1 (conventional; germinal center-inexperienced) and C2 (leukemic non-nodal; germinal center-experienced) MCLs^9^; naïve-like/low-programmed, intermediate/intermediate-programmed and memory-like/high-programmed CLLs^10,11^, and finally germinal center B cell (GCB) and activated B cell (ABC) DLBCLs^32^. In MM, a previous report did not show methylation differences among the distinct cytogenetic subtypes^14^ and thus MM subgrouping was not included in our analyses. Here, we focused on the identification of epigenetic biomarkers that may allow a comprehensive diagnosis of B-cell tumors entities and subtypes. We devised a strategy to construct a classifier algorithm that yielded 56 CpGs as the optimal number distributed along 5 predictors (Extended Data Fig. 3a-e and Supplementary Table 7, Methods) to accurately discriminate the main B-cell tumor entities as a first step (predictor 1), and subsequently B-cell tumor subtypes as a second step (predictors 2, 3, 4 or 5) (Fig. 3a and Methods). The accuracy of the five predictors was evaluated using nested 10-fold stratified cross-validation in the training series (n=1345) and with external validation series (n=711) (Fig. 3b). Overall, we obtained very high accuracies in the predictions in both main B-cell tumor entities (mean sensitivities for training series = 0.97 and validation series =0.99) and B-cell tumor subtypes (mean sensitivities for training series = 0.9 and validation series = 0.97). In some tumor subtypes, we obtained lower accuracy in the predictions mostly due to small sample sizes. The script to easily implement the classifier is provided as supplementary information (SI1). This epigenetic classifier may represent the basis for a simple and accurate diagnostic tool for main B-cell tumors and B-cell tumor subtypes with more complicated diagnosis such as subtypes of MCL or CLL.

**Fig. 3:**
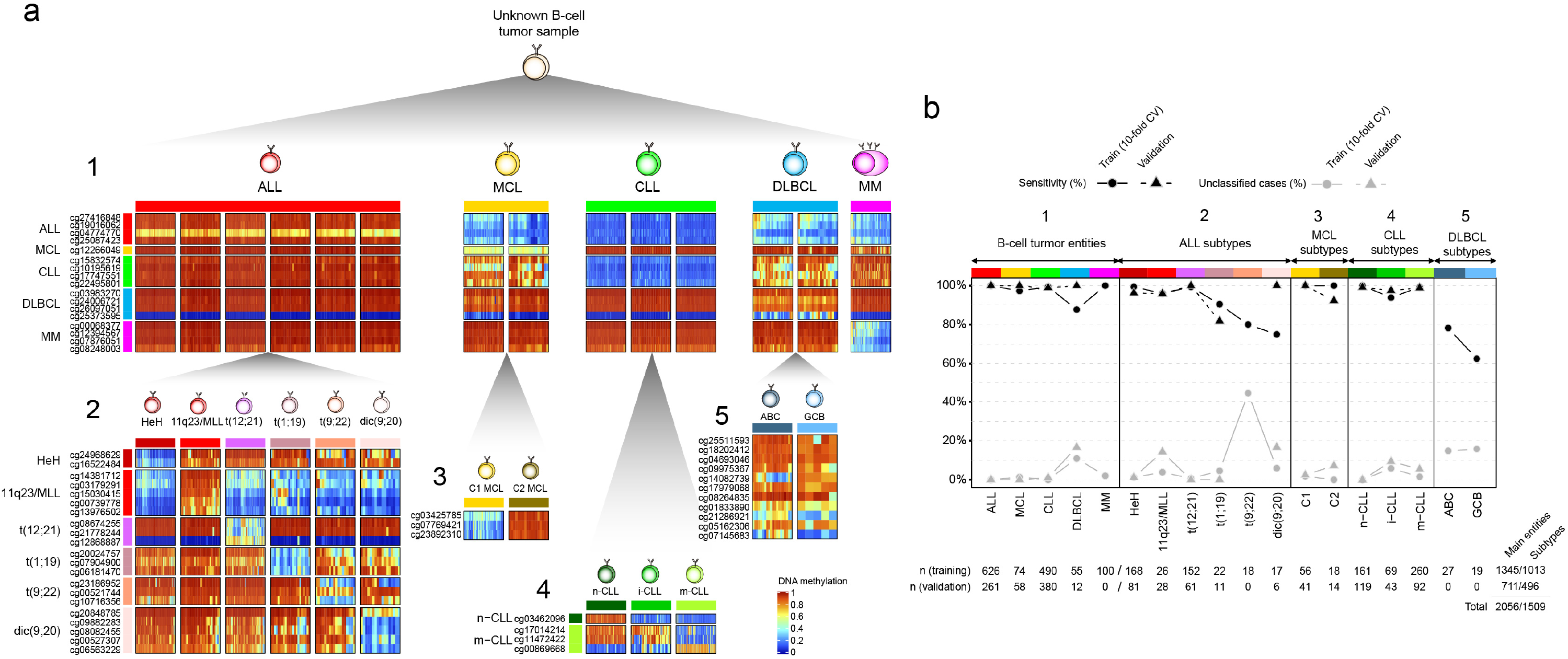
Accurate classification of 14 clinico-biological subtypes of B cell neoplasms using epigenetic biomarkers. **a,** Heatmaps for the CpGs used for the pan B-cell cancer classifier. The classifier consists of two steps: in the first step (1) an unknown B-cell tumor can be predicted into ALL, MCL, CLL, DLBCL or MM, and subsequently (second step, using one of the predictors (2), (3), (4) or (5)) to any B-cell tumor subtype, namely HeH, 11q23/MLL, t(1;19), t(9;22), dic(9;20) for ALL, C1 and C2 for MCL, n-CLL, i-CLL and m-CLL for CLL, and finally GCB or ABC for DLBCLs. ALL, acute lymphoblastic leukemia; MCL, mantle cell lymphoma; CLL, chronic lymphocytic leukemia; DLBCL, Diffuse large B cell lymphoma; MM, multiple myeloma. **b,** Accuracy for the pan-B-cell cancer diagnostic classifier (formed by the 5 predictors in Fig. 3a) for training and validation series. Sensitivity is represented as black circles or triangles for training or validation series, respectively (sensitivity in the training series was evaluated using 10-fold stratified cross-validation). The total number of samples used for both training and validation is shown.

### Patient-specific DNA methylation changes are associated with silent chromatin without an impact on gene expression

After having characterized entity-based sources of DNA methylation variability, we next aimed at studying patient-specific changes within each tumor subtype (Fig. 1b, level 4). To that end, we computed the total number and the number of hyper- and hypomethylation changes in every single patient within each B-cell tumor subtype as compared to HPC (Fig. 4a). As each B-cell tumor entity is derived from a distinct cellular origin, this approach has the advantage of fixing a reference point for all B-cell tumors, and the changes observed can be subsequently dissected into those modulated in normal B-cell maturation and those taking place exclusively in the context of neoplastic transformation (i.e. B-cell independent changes). Overall, we found large differences in the numbers of DNA methylation changes per patient (Fig. 4a and Supplementary Table 8). To analyze whether some B-cell neoplasms show an intrinsically more variable epigenome, we studied the degree of DNA methylation variability in each group, but no differences were observed (Extended Data Fig. 4a).

**Fig. 4:**
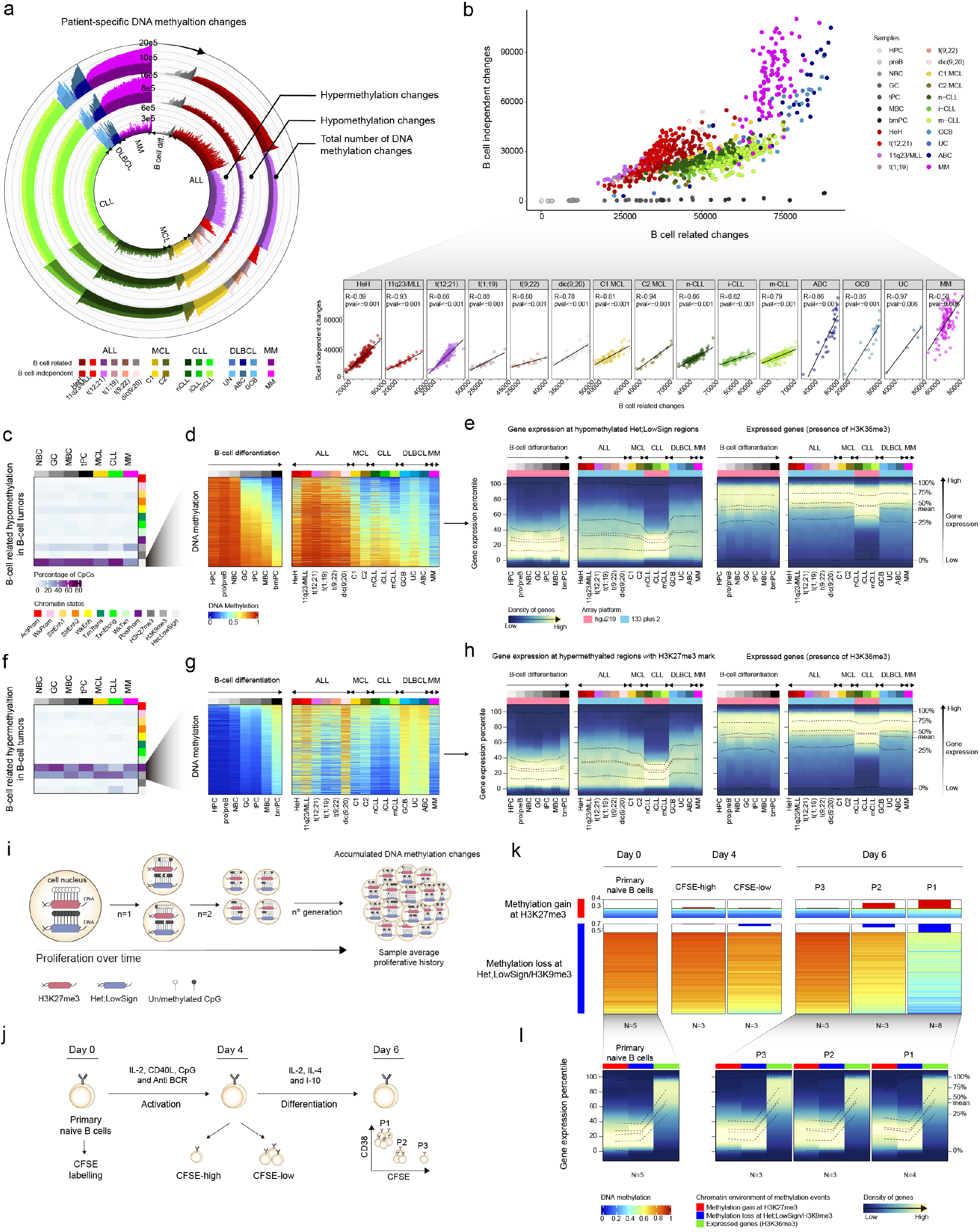
Patient-specific DNA methylation changes are associated with silent chromatin without an impact on gene expression. **a,** Circos plot representing the number of DNA methylation changes with respect to heamatopoietic precursor cells (HPC) in individual patients for normal and neoplastic B cells (each bar represents one patient). Total number of DNA methylation changes, hypomethylation changes and hypermethylation changes are depicted at outer, middle and inner tracks, respectively. Changes are further classified and color-coded as B-cell related or B-cell independent if they occur or not during normal B-cell differentiation, respectively. **b,** Number of B-cell related changes against B-cell independent changes in normal and neoplastic B-cells. Fitted regression lines are shown at bottom per each B-cell tumor subtype. **c,** B-cell related CpGs losing DNA methylation in B-cell tumors and the percentages in each chromatin state in normal and neoplastic B-cells. **d,** Example CpGs from **c** in normal and neoplastic B-cells. **e,** Density of genes distributed along gene expression percentiles of genes associated with B-cell related CpGs losing DNA methylation in B-cell tumors in each B-cell tumor subtype. Expressed genes are displayed at right as control (with presence of H3K36me3). Means within each B-cell subpopulation as well as B-cell tumors are represented. **f,** B-cell related CpGs gaining DNA methylation in B-cell tumors and the percentages in each chromatin state in normal and neoplastic B-cells. **g,** Example CpGs from **f** in normal and neoplastic B-cells. **h,** Density of genes distributed along gene expression percentiles of genes associated with B-cell related CpGs gaining DNA methylation in B-cell tumors in each B-cell tumor subtype. Expressed genes are displayed at right as control (with presence of H3K36me3). Means within each B-cell subpopulation as well as B-cell tumors are represented. **i,** Model for DNA methylation changes occurring at repressed regions during mitotic cell division. **j,** *In vitro* model for plasma blast differentiation from human primary naïve B cells. Primary naïve B cells are labeled with CFSE at day 0. DNA methylation profiles are obtained at day 0 (primary naïve B cells) day 4 (CFSE-high and CFSE-low) and day 6 (P1-CFSE-low and CD38+, P2-CFSE-intermediate, CD38-, P3-CFSE-high CD38-). **k,** DNA methylation changes accumulate at repressed regions during mitotic cell division upon *in vitro* proliferation and differentiation of primary naïve B cells to plasma blasts. Hypermethylation takes place at H3K27me3 regions, whereas hypomethylation at heterochromatin and H3K9me3 regions. ChIP-seq data for primary NBC and GC B cells was used. The mean of several biological replicates is represented (numbers are depicted at bottom of the heatmaps). **l,** Density of genes distributed in gene expression percentiles for genes with changes in DNA methylation in (K).

The total number of altered CpGs per case was classified into four categories, depending whether DNA methylation was gained or lost and whether it was modulated or not during normal B cell development^6^ (Fig. 2b and Fig. 4a). Regardless of the cellular origin of each B-cell tumor entity, this analysis uncovered striking correlations between the degree of B-cell related and B-cell independent DNA methylation changes, a finding that was maintained when hypermethylation and hypomethylation were studied separately (Fig. 4b and Extended Data Fig. 4b). This association suggests that the overall DNA methylation burden of the tumor in each individual patient may be shaped by a similar underlying phenomenon, which is manifested in all four CpG categories. This statement is further supported by a thorough annotation of the CpGs sites. Patient-specific CpGs that undergo hypomethylation in the B-cell related and B-cell independent fractions are consistently located in low CpG-content (open sea) low-signal heterochromatin, and the associated genes are constitutively silent both in normal and neoplastic B cells (Fig. 4c-e and Extended Data Fig. 4c-f). In the case of patient-specific hypermethylation, CpGs in both fractions are located in promoter regions and CpG islands (CGIs) with H3K27me3-repressed and poised-promoter chromatin states, and affect genes that remain silent across normal differentiation and neoplastic transformation of B cells (Fig. 4f-h, Extended Data Fig. 4c, g-i). These findings indicate that most DNA methylation changes in B-cell tumor patients occur in silent chromatin regions in the absence of concurrent phenotypic changes, suggesting that a mechanism independent from gene regulation may underlie their overall DNA methylation landscape.

Beyond the classical role of DNA methylation as gene regulator, an accumulated body of published evidence supports the concept that DNA methylation changes accumulate during cell division (Fig. 4i)^33–39^. Studies in fibroblasts and hematopoietic stem cells have reported that DNA methylation loss in late-replication heterochromatic regions and DNA methylation gain of polycomb targets increase as cells proliferate without an apparent impact on gene expression^33–35^. Furthermore, DNA methylation loss in heterochromatic, late-replicating domains has been recently associated with mitotic cell division in cancer^39^. Additionally, early studies detected preferential methylation of CGI marked by H3K27me3 in cancer^40–42^, and DNA methylation changes in regions marked by H3K27me3 in embryonic stem cells have been recently related to mitotic cell division in cancer^38^.

In order to explore whether DNA methylation changes in silent regions reflect the proliferative history of primary human B cells, we used DNA methylation data of an *in vitro* differentiation model of primary NBCs into plasma cells (Fig. 4j)^43^. At days 4 and 6, different B cells were separated based on their proliferation history measured by CFSE dilution. We evaluated the DNA methylation profile in repressed regions at these time points and we detected the presence of hypermethylation at repressed H3K27me3-containing regions and hypomethyation of low signal/H3K9me3-containing heterochomatic regions in cells that have proliferated (Fig. 4k). This finding was particularly marked at day 6, in which the gradual accumulation of DNA methylation changes in silent regions was directly associated with their proliferative history (from less divided P3 to highly divided P1). The genes associated to the measured CpGs did not show any change of expression levels regardless of their methylation status, and thus were unlikely to be related to the phenotype of the cells (Fig. 4l).

Collectively, all these data indicate that a great fraction of the patient-specific DNA methylation changes in B-cell tumors accumulate at H3K27me3 and low signal/H3K9me3 regions during cell division without affecting gene expression levels, a finding that is consistent with an epigenetic mitotic clock.

### Development and validation of an epigenetic mitotic clock reflecting the proliferative history of normal and neoplastic B cells

We performed a step-wise selection of CpGs whose methylation status would reflect the cell mitotic activity regardless of its normal or neoplastic nature (Methods). Using this strategy, we finally retained 184 CpGs in stable polycomb regions that tend to gain DNA methylation and 1,164 CpGs in constitutive heterochromatin that tend to lose DNA methylation during normal B-cell maturation and neoplastic transformation (Fig. 5a, Supplementary Table 9 and SI2). We next constructed two scores, one for hypermethylation and one for hypomethylation, which we respectively named epiCMIT-hyper and epiCMIT-hypo, ranging from 0 to 1 depending on low or high proliferative histories, respectively (Methods). Remarkably, epiCMIT-hypo contained a significant proportion of solo-WCGW CpGs, which have been recently associated with human mitotic cell division (N=1e5 permutations, pval<0.0001)^39^. As normal B cell differentiation entails cell division, we initially evaluated both scores in normal B cells and observed an expected but strikingly high correlation (R=0.96, pval<0.001), with B-cell subpopulations distributed according to their maturation state (and thus according to their accumulated proliferative history during B-cell differentiation) (Fig. 5b, left panel). This finding suggests that mitotic cell division in normal B cells leaves both hyper- and hypomethylated imprints. This high correlation between the two scores was also observed for MCL, CLL and DLBCL (Fig. 5b) but not for ALL and MM. The epiCMIT-hyper was greater than the epiCMIT-hypo in ALL samples, and the opposite scenario was observed in MM. As these two neoplasias originate from B cells at the two extremes of the maturation spectrum, our data suggests that the impact of cell division onto the DNA methylome may be different depending on the maturation state of the cellular origin. Upon neoplastic transformation, dividing precursor B cells do not seem to acquire broad hypomethylation in heterochromatin but rather hypermethylation in polycomb-repressed regions. In contrast, neoplastic plasma cells acquire widespread hypomethylation in heterochromatin and virtually lack hypermethylation of polycomb-repressed regions. As previously shown in Extended Data Fig. 2d, this phenomenon may be related to the differential expression of DNMTs of the respective cells of origin of ALL and MM. We then took into consideration both scores to derive a unique epiCMIT score (Fig. 5c). The epiCMIT reflects the relative accumulation of mitotic cell divisions of a particular sample and is able to capture different tendencies in gaining or losing DNA methylation during mitotic cell division, as it respectively happens in ALL and MM patients (Fig. 5b). Finally, epiCMIT cannot be affected by different distribution of cell cycle phases in tumor samples, since the DNA methylome remains rather stable during the whole cell cycle^44^.

**Fig. 5:**
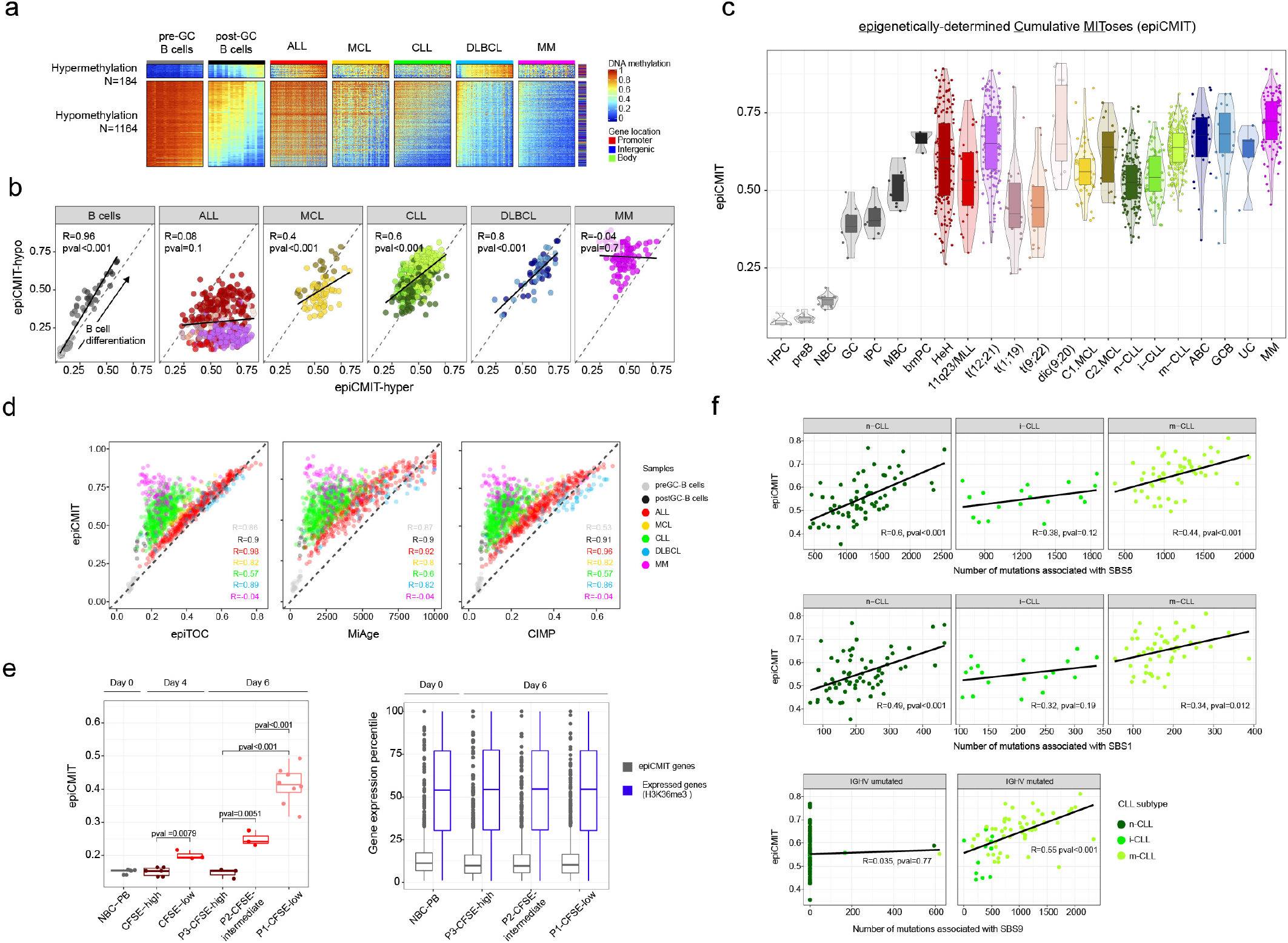
Development and validation of an epigenetic mitotic clock reflecting the proliferative history of normal and neoplastic B cells. **a,** Selection of CpGs gaining DNA methylation at H3K27me3 and losing DNA methylation in normal and neoplastic B cells to construct the epiCMIT-hyper and epiCMIT-hypo scores. epiCMIT, epigenetically-determined Cumulative MIToses. **b,** Correlation of epiCMIT-hyper and epiCMIT-hypo in normal and neoplastic B cells. **c,** epiCMIT in normal and neoplastic B cells. **d,** Correlation of epiCMIT with previously reported hypermethylation-based mitotic clocks, including epiTOC (epigenetic Timer for Cancer risk) and MiAge. Correlation with CIMP (CpG hypermethylator phenotype) is also shown, although it has not been formally presented as mitotic clock. **e,** epiCMIT score during the *in vitro* B-cell differentiation as well as gene expression of genes containing epiCMIT-CpGs. Expressed genes are also shown as control (presence of H3K36me3). **f,** Correlation of epiCMIT and mutational signatures related to proliferative history for 138 CLL samples with available DNA methylation data and WGS. SBS1, SBS5 have been reported as clock-like mutational signatures. SBS9 is related to ncAID mutations.

The applicability of the epiCMIT as mitotic clock was validated through several perspectives. First, we compared it with two previously reported hypermethylation-based mitotic clocks called epiTOC and MiAge^38,45^ (Supplementary Table 8). The epiCMIT showed excellent correlations with these clocks in B-cell neoplasms that tend to acquire polycomb-related hypermethylation (e.g. mostly ALL, but also DLBCL and MCL); a moderate correlation in the case of CLL, which acquires more hypo-than hypermethylation (Fig. 4a and Extended Data Fig. 4b), and a total lack of correlation in the case of MM, which mostly loses DNA methylation (Fig. 5d and Extended Data Fig. 5a). Identical observations were obtained comparing the epiCMIT and the widely-reported CpG island methylator phenotype (CIMP) in human cancer^46^ calculated as previously proposed^47^ (Supplementary Table 8). Therefore, our analysis also reveals that the classical CIMP score in reality may represent a measure of the mitotic history of the cells. A potentially confusing aspect related to epiCMIT is the fact that DNA methylation has mainly been used to measure the chronological age of an individual^48,49^. In order to study the relationship between mitotic and aging DNA methylation clocks, we first compared the epiCMIT in naïve B cells from infant, adults and elderly donors and showed that epiCMIT was stable in all age ranges regardless of donor’s age (Extended Data Fig. 5b). We next applied the most popular chronological age clock proposed by Horvath^50^ and showed that the age of donors was predicted with high accuracy. This result indicates that, although both Horvath and epiCMIT scores are based on DNA methylation, they measure distinct biological phenomena (Extended Data Fig. 5b). Furthermore, the epiCMIT was highly variable in pediatric ALL samples with a minimum age range (Extended Data Fig. 5c). Collectively, these analyses indicate that the epiCMIT is a more universal mitotic clock than hypermethylation-based clocks since it captures the proliferative history of neoplasms regardless of their tendency of gaining or losing DNA methylation during mitotic cell division.

Second, the epiCMIT was validated using the *in vitro* B-cell differentiation model shown in Fig. 4j, in which we observed that epiCMIT increases with the proliferative history of the cells without altering gene expression levels of the epiCMIT-associated genes (Fig. 5e). Third, using WGS data from 138 CLL patients from our cohort^10,17,51,52^, we observed that the epiCMIT was highly correlated with the total number of somatic mutations (Extended Data Fig. 5d). We next extracted mutational signatures as recently described^53^ (Extended Data Fig. 5e) and observed significant correlations with mutational signatures SBS1 and SBS5 (Fig. 5f), that have been recently described as mitotic-like mutational processes^54^. We also identified a significant link between the epiCMIT and the non-canonical AID signature (SBS9) (Fig. 5f)^52,55^ in IGHV mutated CLL, possibly reflecting rounds of cell divisions in germinal center B cells before differentiation into memory B cells and malignant transformation. Fourth, although the epiCMIT reflects the proliferative history of the cell rather than the actual proliferative status of the samples (e.g, bmPC have high epiCMIT and do not proliferate, Fig. 5c), a relationship between epiCMIT and cell proliferation is expected (more proliferative history implies overall more proliferation, although also depends on time). Accordingly, MCL cases showing higher Ki-67 (a proliferation marker) also had higher epiCMIT than cases with moderate Ki-67 expression (Extended Data Fig 5f). Furthermore, GSEA analyses in ALL and CLL cases with high and low epiCMIT revealed that cases with high epiCMIT showed higher expression of genes related with cell proliferation (Extended Data Fig. 5g, h). Thus, these data suggest that cases with higher proliferative history also seem to have higher proliferation at the time of sampling. Collectively, the four lines of evidence presented above support that the epiCMIT may represent a *bona fide* measure of the relative number of cumulative mitotic cell divisions that normal and neoplastic B cells have undergone since the uncommitted hematopoietic cell stage.

### The epiCMIT is a strong independent variable predicting clinical behavior in B-cell tumors

In normal B-cell maturation, the epiCMIT gradually augments as B cells proliferate during cell differentiation, an increase that is particularly marked in proliferative GC B cells (Fig. 5c). In neoplastic B cells, however, the interpretation of the epiCMIT is less trivial and must be divided into two components: the epiCMIT of the cell of origin and the epiCMIT acquired in the course of the neoplastic transformation and progression (Extended Data Fig. 6a). Therefore, the relative epiCMIT must be compared among patients from entities arising from the same B-cell maturation stage and should be a dynamic variable during cancer progression. Thus, we compared the epiCMIT in two paradigmatic transitions between precursor conditions and overt cancer, i.e. monoclonal gammopathy of undetermined significance (MGUS) and MM, as well as monoclonal B cell lymphocytosis (MBL) and CLL categorized according to their cellular origin. This analysis showed an overall lower epiCMIT in precursor lesions compared with overt cancer (Fig. 6a, upper panels), as would be expected due to their increased leukemia-specific proliferative history. In line with this finding, the epiCMIT increased in paired CLLs at diagnosis and progression before treatment as well as in sequential ALL samples at diagnosis, first relapse and second relapse (Fig. 6a, lower panels). These results suggest that the epiCMIT evolves together with clinical progression.

Based on our previous observations, we next wondered whether the epiCMIT could be useful to predict the clinical behavior of B-cell neoplasms. We analyzed the relationship of the epiCMIT with clinical outcome in specific B-cell tumor subtypes based on their shared cytogenetics (i.e. ALL) or cell of origin (i.e. MCL, CLL and DLBCL), and thus having a similar ground state B-cell specific proliferative history (Extended Data Fig. 6a). In ALL, high epiCMIT was consistently associated with longer relapse-free and overall survival (Fig. 6b and Extended Data Fig. 6b) of the patients within each cytogenetic subtype, and thus better clinical outcome. Next, we additionally showed epiCMIT as significant variable in a multivariate regression Cox model with epiCMIT as continuous variable together with cytogenetic subtypes (Fig. 6b). This observation is in line with published evidence showing that a higher CIMP, which is highly correlated with the epiCMIT in ALL (Fig. 5d), is a good prognosis factor in this disease^56,57^. In contrast to ALL, the opposite clinical scenario was observed in mature B-cell neoplasms. In each of the CLL subtypes, a high epiCMIT was strongly associated with a worse prognosis using time to treatment (TTT) as end-point variable, both from sampling time (Fig. 6c, left panel) and in cases whose sample was obtained close to diagnosis (Extended Data Fig. 6c, left panel). Additionally, a multivariate Cox regression model for TTT revealed that epiCMIT as continuous variable was a highly significant variable conferring dismal prognosis together with age, number of driver genetic alterations^17,58^ and epigenetic subgroups^10,11,59^ (Fig. 6c right panel and Extened Data Fig. 6c). The epiCMIT maintained its independent prognostic value using overall survival as end-point variable, although its effect was moderate (Extended Data Fig. 6d). These findings were widely confirmed in an additional series of 136 CLLs treated with chemo-immunotherapy (Fig. 6d and Extended Data Fig. 6c, d right panels). In the case of MCL, the epiCMIT showed an independent poor prognostic impact in the two cell-of-origin subtypes (C1 and C2), an observation that was confirmed in an extended series in the more aggressive and prevalent C1 group (Fig. 6e, f). Lastly, although the sample size was limited and requires further studies, our data suggest that high epiCMIT could also represent a poor prognostic variable within the two cell-of-origin DLBCL subtypes (Extended Data Fig. 6e). Overall, these results suggest that an increased proliferative history of the neoplastic clone in precursor B-cell neoplasms at diagnosis predicts for a better disease-free survival, as previously published in studies analyzing the CIMP score^56,57^. In sharp contrast, in mature B-cell neoplasms, which are overall less proliferative than ALL, neoplastic clones with high proliferation history seem to predict for future proliferative capacity and consistently show worse clinical outcomes.

### epiCMIT is associated with specific genetic driver alterations in CLL

We next sought to assess which CLL driver alterations may confer a proliferative advantage to neoplastic cells, and subsequently a higher epiCMIT. To that end, we exploited 477 CLL samples in which we had DNA methylation data and whole exome sequencing (WES) (Fig. 7a). We initially depicted all driver genetic changes in each CLL subtype divided in high and low epiCMIT (Extended Data Fig. 7a). Next, we interrogated the levels of epiCMIT in patients with each driver genetic alteration (with at least 4 mutated patients). We performed the analyses in all CLL patients (showed as Global in Fig. 7b) and then within each epigenetic subgroup separately (Fig. 7b, Extended Data Fig. 7b and Methods). We showed significant and positive associations of epiCMIT with 24 genetic driver alterations affecting the main signaling pathways altered in CLL (Fig. 7b,c)^17,58^. The majority of these genetic alterations have been previously linked to an adverse clinical behavior of patients, such as *NOTCH1, TP53, SF3B1, ATM, BIRC3 or EGR2.* Interestingly, epiCMIT showed a clear association with a new and recently identified non-coding genetic driver in CLL, the U1 spliceosomal RNA, a finding that may help in explaining its suggested poor prognostic impact^60^.

**Fig. 6:**
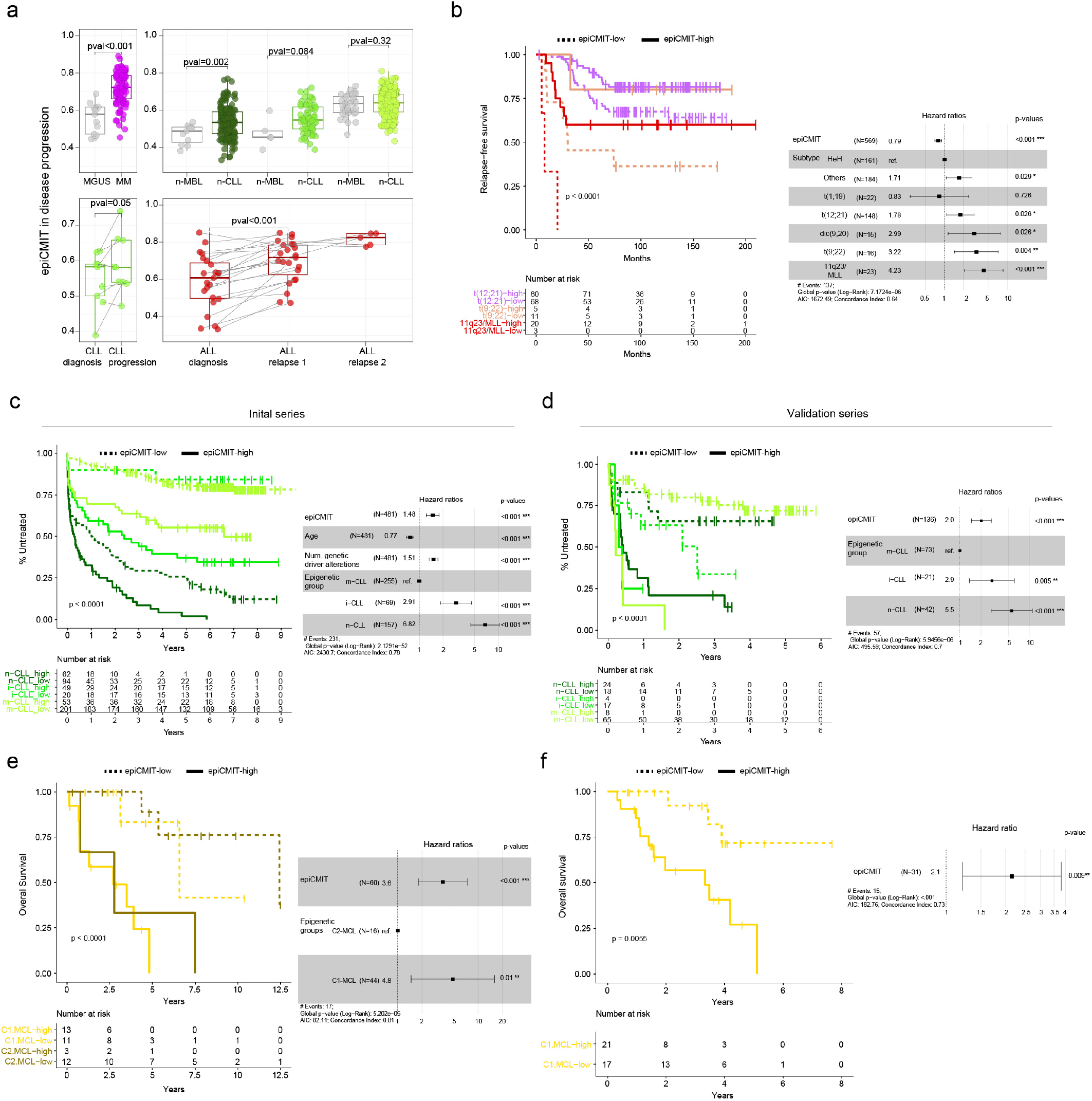
The epiCMIT is a strong independent variable predicting clinical behavior in B-cell tumors. **a,** epiCMIT is a dynamic variable that reflects disease progression. Precursor conditions MGUS and MBL show significantly lower epiCMIT levels than their respective cancer conditions CLL and MM, respectively. epiCMIT increases from diagnosis to progression in paired CLL samples and in paired ALL at diagnosis, first relapse and second relapse. **b,** Kaplan-Meier curves for three ALL cytogenetic groups shown as example divided in low and high epiCMIT using maxstat rank statistics. Multivariate cox regression model for relapse-free survival with epiCMIT against cytogenetic groups. Hazard ratio for epiCMIT correspond to 0.1 increments. **c,** Kaplan-Meier curves for CLL epigenetic groups based on different cellular origin divided in low and high epiCMIT using maxstat rank statistics. Multivariate cox regression model for time to first treatment with epiCMIT against age, number of driver alterations and epigenetic groups based on different cellular origin. Validation series is shown in **d**. Hazard ratio for epiCMIT correspond to 0.1 increments. **e,** Kaplan-Meier curves for MCL epigenetic groups based on different cellular origin divided in low and high epiCMIT using maxstat rank statistics. Multivariate cox regression model for overall survival with epiCMIT against epigenetic groups based on different cellular origin. Hazard ratio for epiCMIT correspond to 0.1 increments. **f,** Validation series for C1 MCL. Hazard ratio for epiCMIT correspond to 0.1 increments.

Remarkably, the presence of some genetic alterations was associated with high epiCMIT indistinctly in all patients, such as *TP53*, while others were particularly associated with epiCMIT within CLL subgroups, such as *NOTCH1* in n-CLL and i-CLL, *SF3B1* in i-CLL, and del(13q) and tri12 in m-CLL.

Collectively, these results suggest that the well-established clinical impact of genetic alterations in CLL may be explained by their association with a high proliferative potential, being this association different for certain genetic alterations depending on the maturation state of the cellular origin.

## Discussion

In this study, we have followed a systematic approach to dissect the sources of DNA methylation variability of B-cell neoplasms in the context of the normal B-cell differentiation program. Overall, we found that the methylation levels of 88% of all CpGs are modulated in normal and/or neoplastic B cells, suggesting that the human DNA methylome is even more dynamic than previously appreciated^6,61,62^. The extensive DNA methylation variability among different B-cell neoplasms is in part related to imprints of normal B-cell development. This phenomenon has been previously observed and has led to a more accurate classification not only of B-cell neoplasms^9–11,59^, but also of solid tumors^2,63,64^. In addition to this epigenetic link to normal cell maturation, each B-cell neoplasm also shows disease-specific hyper- and hypomethylation. Of particular interest are the disease-specific *de novo* hypomethylation signatures in active regulatory regions, which were associated with binding of TFs involved in the pathogenesis of each respective B-cell tumor. This phenomenon was particularly marked in ALL, MCL and CLL, whose *de novo* hypomethylation was enriched in regulatory elements^23–25^. Unexpectedly, although DLBCL and MM pathogenesis has been linked to TFs, *de novo* hypomethylation was depleted in regulatory elements containing TF binding sites. In these two malignancies, we detected few binding sites of TFs with potential involvement in the diseases. However, we did not detect classical TFs such as BCL6 in DLBCL or IRF4 in MM, possibly because they are key players during B-cell differentiation and their binding sites may already be hypomethylated in the normal B-cell counterparts of DLBCL and MM.

In spite of the importance of DNA methylation in regulatory regions, we identified that the majority of DNA methylation changes in B-cell neoplasms, and especially those related to inter-patient variability within each B-cell tumor subtype, are located in inactive chromatin. The magnitude of this inter-patient variability affecting normal B-cell related and -independent regions is highly correlated, suggesting that the cause of these DNA methylation changes is a phenomenon that takes place both in normal and neoplastic B cells. Compelling published evidence as well as new data presented in our study support the notion that mitotic cell division leaves transcriptionally-inert epigenetic imprints onto the DNA. These take place in the form of hypomethylation of heterochromatin and hypermethylation of polycomb-repressed regions. This knowledge has recently led to the concept of using DNA methylation as a mitotic clock^38,45^, and has been recently used by single cell DNA methylation data to track the evolutionary trajectory of CLL^65^. Notably, methylation-based mitotic clocks seem to capture different biological information from aging clocks (Extended Data Fig. 5b)^48^. As far as we are aware, there are two published mitotic clocks^38,45^ and both are hypermethylation-based. Furthermore, our analyses suggest that the classical CIMP phenomenon in cancer^46^ may indeed represent another hypermethylation-based mitotic clock. However, only using hypermethylation to determine the mitotic clock is insufficient to capture the mitotic activity of the cells, as some neoplasms may not acquire hypermethylation upon cell division but rather hypomethylation. For instance, neoplastic plasma cells in MM do not tend to acquire polycomb-related hypermethylation. Thus, using hypermethylation to determine the mitotic history of MM cells would incongruently lead to the conclusion that they have not proliferated beyond their cellular origin (Fig. 5d and Extended Data Fig. 5a). Therefore, to circumvent possible misleading interpretations, we generated a more broadly applicable mitotic clock (called epiCMIT) that takes into consideration both hyper- and hypomethylation. Importantly, epiCMIT aims at capturing the entire mitotic history of cells, including cell division associated both with normal development as well as neoplastic transformation and progression (Extended Data Fig. 6a). Thus, the epiCMIT must not be compared among tumors arising from different normal counterparts but its relative magnitude must be studied in those arising from a particular maturation stage. Within each of these subgroups, the relative epiCMIT has a profound independent prognostic value from other well-established clinical variables. Increased epiCMIT is associated with worse clinical outcome within CLL and MCL subgroups, thus indicating that superior proliferative history before treatment seems to determine future proliferative capacity and is thus associated with worse clinical outcome. Strikingly, we consistently found the opposite pattern in precursor ALL subgroups, a finding in line with recent reports showing that the presence of CIMP (a hypermethylation-based mitotic clock) is associated with better clinical outcome^56,57^. These results suggest that children having more proliferative ALL cells at diagnosis (and thus a larger proliferative history) are more prone to be cured with high intensive chemotherapy regimens,^66^, which cannot be administrated in elderly patients.

Finally, in order to identify the genetic lesions associated with higher proliferative history in CLL, we exploited our thorough genetic characterization of our cases to study the relationship between driver genetic alterations and epiCMIT. We identified 24 driver genetic alterations that may confer a higher proliferative capacity and thus are associated with higher epiCMIT or methylation evolution^67^. These genetic alterations were distributed throughout the main altered signaling pathways in CLL and were manifested differently in distinct CLL subgroups based on their cellular origin (Fig. 7b, c). This finding suggests that specific mutations may predispose to a higher proliferative advantage depending on the maturation stage and (epi)genetic makeup of the cellular origin.

**Fig. 7:**
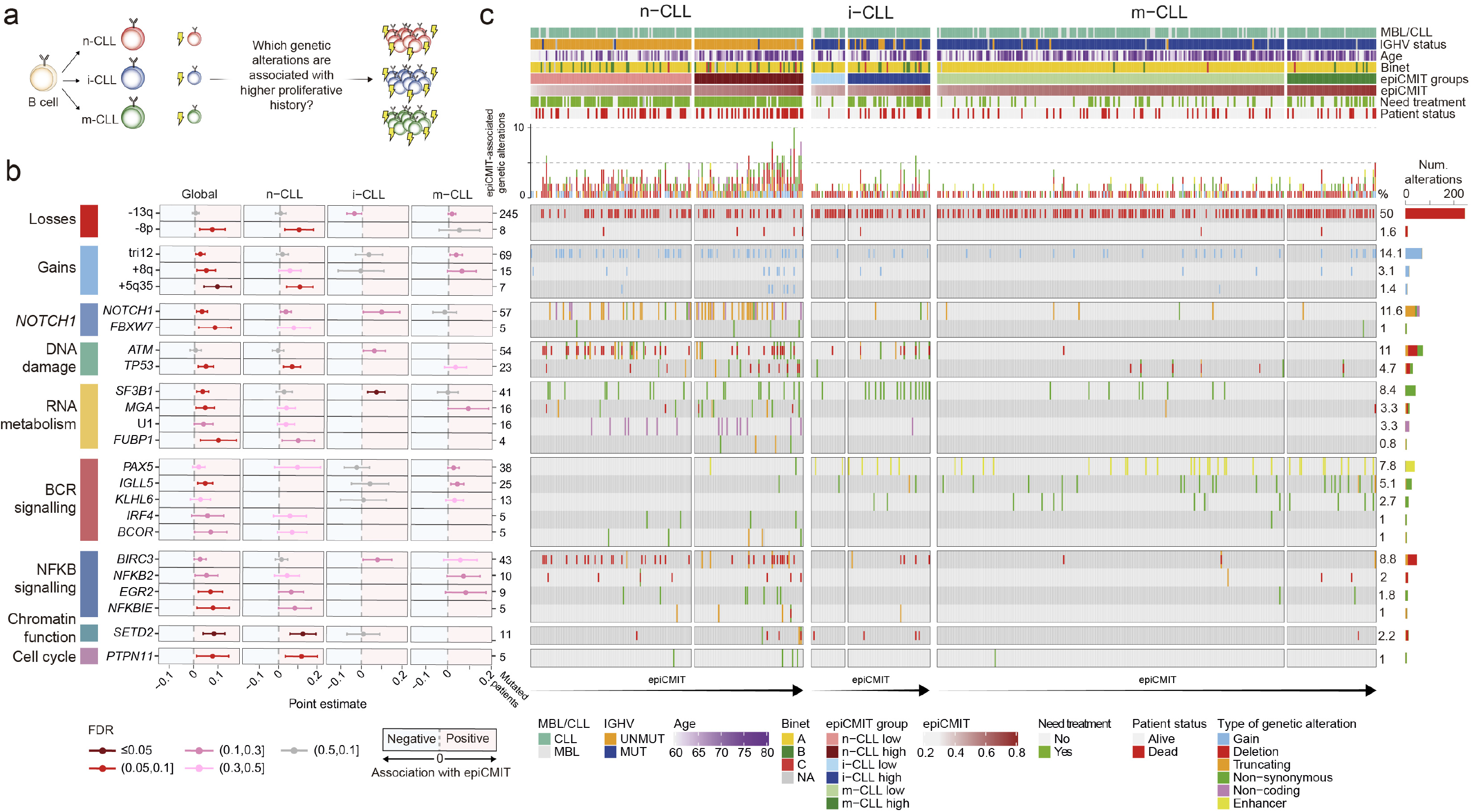
epiCMIT is associated with specific genetic driver alterations in CLL. **a,** Are there specific genetic alterations in CLL that may predispose a proliferative advantage to neoplastic cells? **b,** Genetic driver alterations in CLL associated with higher epiCMIT grouped by signaling pathways reported. Analyses were done globally for all CLL samples (although adjusted by epigenetic groups) as well as within each epigenetic subgroup. Positive point estimates relate to positive associations with epiCMIT. 95% confidence intervals are shown with colors coding for FDR corrections. Number of patients with each genetic driver alteration is shown at right. **c,** Oncoprint showing genetic driver alterations associated with higher epiCMIT in CLL epigenetic groups separately. Other clinicobiological features including MBL or CLL, IGHV status, Age, Binet stage, epiCMIT subgroups based on Fig. 5c, need for treatment and patient status are shown. Cases are ordered within each CLL subgroup from lower to higher epiCMIT values. Distinct genetic driver alterations are depicted with different colors and shapes. The percentage of mutated patients as well as barplots showing the number of mutated patients for each alteration is shown at right.

In summary, our comprehensive epigenetic evaluation of normal and neoplastic B cells at different maturation stages uncovers multiple new insights into the biological roles of DNA methylation in cancer, an analytic approach that may also benefit our understanding of other cancers. From a clinical perspective, DNA methylation may provide a holistic diagnostic and prognostic approach to B-cell neoplasms. Particularly, we defined an accurate and easy-to-implement pan-B-cell tumor diagnostic tool and generated a mitotic clock reflecting the proliferative history of the neoplastic cells of each patient to estimate their clinical risk, which shall represent a valuable asset in the precision medicine era.

## ACKNOWLEDGEMENTS

This research was funded by the European Union’s Seventh Framework Programme through the Blueprint Consortium (grant agreement 282510), Generalitat de Catalunya Suport Grups de Recerca AGAUR 2017-SGR-1142 (to E.C.) and 2017-SGR-736 (to J.I.M.-S.), Ministerio de Ciencia, Innovación y Universidades of the Spanish Government (MCIU), Grants RTI2018-094274-B-I00 (to E.C.) and SAF2017-86126-R (to J.I.M.-S.) as well as Proyecto Medicina Personalizada PERMED (Grant PMP15/00007), which is part of Plan Nacional de I+D+I and is co-financed by the ISCIII-Sub-Directorate General for Evaluation and the European Regional Development Fund (FEDER-“Una manera de Hacer Europa”), CIBERONC (CB16/12/00225, CB16/12/00334, CB16/12/00236, and CB16/12/00489), the Accelerator award CRUK/AIRC/AECC joint funder-partnership, research funding from Fondo de Investigaciones Sanitarias, Instituto de Salud Carlos III PI17/01061 (SB), Ministerio de Ciencia, Innovación y Universidades (MCIU), RTI2018-094274-B-I00, SAF2015-64885-R (EC), the NIH grant number 1 P01CA229100 (EC), and the European Regional Development Fund “Una manera de fer Europa”, CERCA Programme/Generalitat de Catalunya. FN is supported by a pre-doctoral fellowship of the Ministerio de Economía y Competitividad (MINECO, BES-2016-076372). E.C. is an Academia Researcher of the “Institució Catalana de Recerca i Estudis Avançats” (ICREA) of the Generalitat de Catalunya. This work was partially developed at the Centro Esther Koplowitz (CEK, Barcelona, Spain). We thank Francesco Maura for his help with the analysis of mutational signatures.

## AUTHOR CONTRIBUTIONS

Investigator contributions were as follows: T.B., J.N., Y.N-Z., G.L., A.R-D., S.M., R.O., G.C., M.K., A.C-Q., L.S-T., J.W., J.L., E.G., S.B., P.J., X.A., F.P., C.L-O., X.S.P., C.C.O., T.Z., J.D., A.L-G. and E.C. contributed to sample biological and/or clinical annotation; M.D-F., G.C., F.N. and R.B. performed methylome, ChIP-Seq, transcriptome, genetic and/or statistical analyses; R.R., M.P. and D.T. provided computational support. M.D-F. and J.I.M.-S. participated in the study design. M.D-F., F.N., R.B., T.B., J.D., A.L-G., E.C. and J.I.M.-S. participated in data interpretation. J.I.M.-S. directed the research and wrote the manuscript together with M.D-F.

## sCOMPETING INTERESTS

The authors declare no competing interests.

## Supplementary figure legends

**Extended Data Fig.1.**
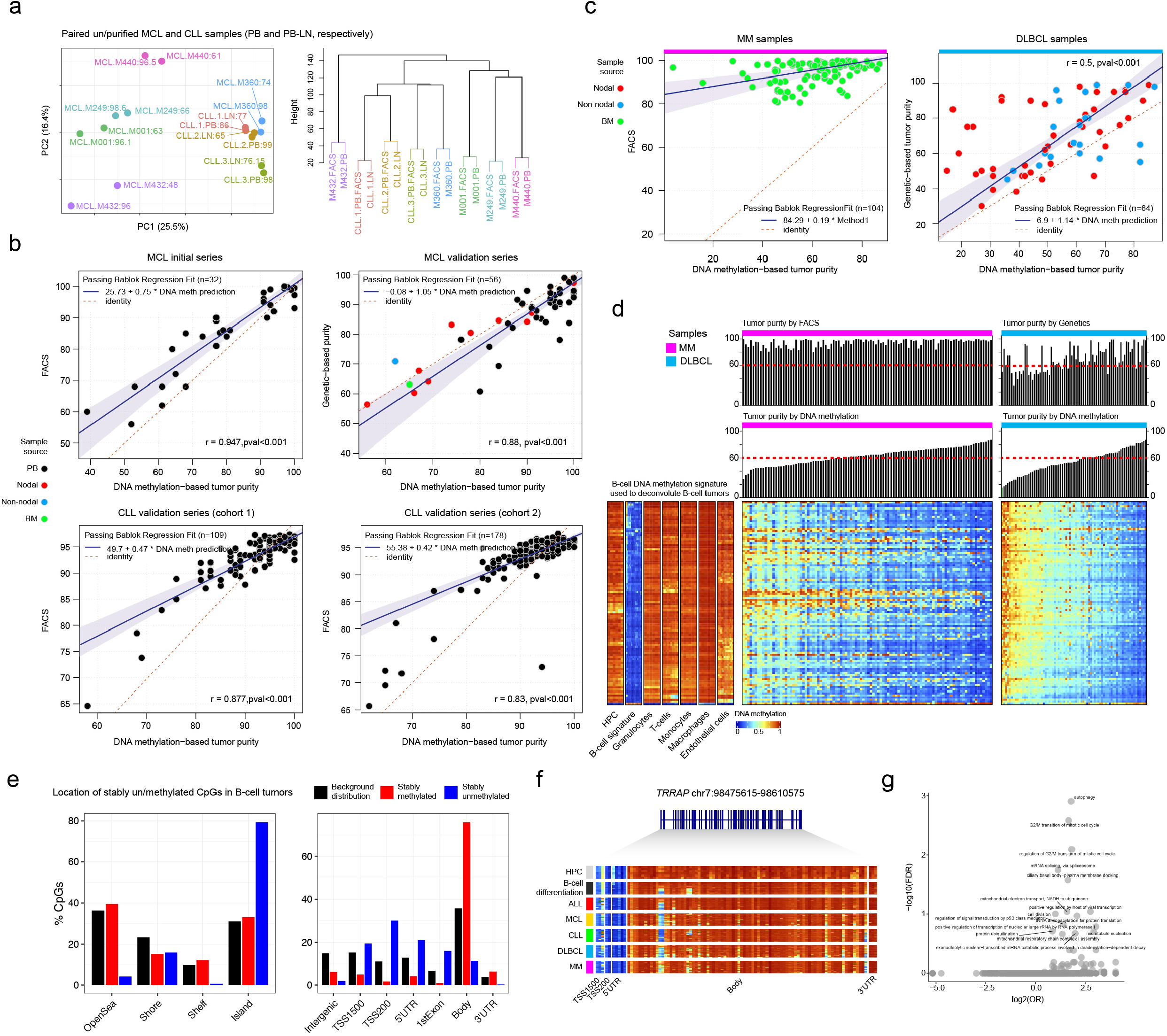
**a,** Principal component analysis and hierarchical clustering of paired un/purified DNA methylation profiles obtained with EPIC array from MCL and CLL patients. Colors represent the same un/purified sample, with FACS-based purities highlighted in each sample. MCL, mantle cell lymphoma. CLL, chronic lymphocytic leukemia. **b,** Correlations and Passing Bablock regression fits of gold-standard methods for tumor purity prediction (FACS and genetic-based) against DNA methylation-based tumor purity prediction for MCL and CLL patients. **c,** Pearson correlations and Passing Bablock regression fits for gold-standard methods for tumor purity predictions (FACS and genetic-based) against DNA methylation-based tumor purity predictions for MM and DLBCL patients. MM, multiple myeloma. DLBCL, Diffuse large B cell lymphoma. **d,** Pan-B cell DNA methylation signature used to deconvolute DNA methylation and obtain B-cell tumor purities in DLBCL and MM. Bar plots representing DNA-methylation based predictions as well as gold standard-based predictions are represented at top the heatmaps. **e,** Genomic distribution of stably un/methylated CpGs in B-cell tumors. **f,** Example gene showing stably un/methylated CpGs. **g,** Gene ontology analysis of genes showing both stable un/methylated CpGs.

**Extended Data Fig.2.**
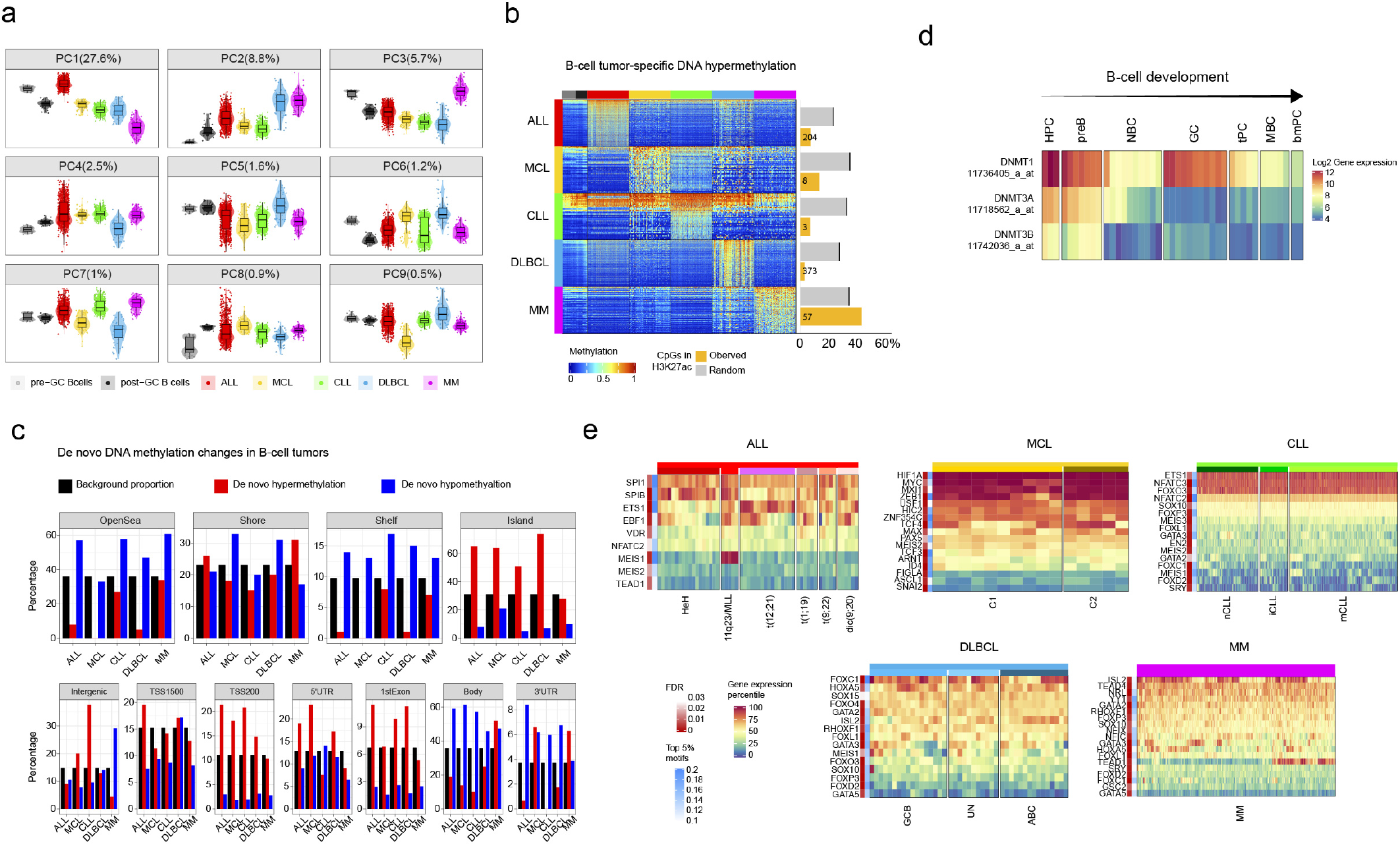
**a,** Principal component analysis for normal and neoplastic B cells, including ALL, MCL, CLL, DLBCL and MM. Each component is resented separately (until the ninth component). ALL, acute lymphoblastic leukemia; MCL, mantle cell lymphoma; CLL, chronic lymphocytic leukemia; DLBCL, Diffuse large B cell lymphoma; MM, multiple myeloma. **b,** Heatmap showing B-cell tumor-specific hypermethylation and the number of CpGs falling at regulatory regions. Per each B-cell tumor, the same number of *de novo* CpGs was randomly chosen from the 450K array and interrogated the percentage falling at regulatory regions. **c,** Genomic distribution for *de novo* DNA methylation changes in B-cell tumors. **d,** Gene expression of DNA methyltransferases *DNMT1, DNMT3A* and *DNMT3B* throughout B cell differentiation. **e,** Gene expression percentile of TFs showing the most significant p-values and frequencies in *de novo* hypomethylation signatures in each B-cell tumor in Fig. 2d.

**Extended Data Fig.3.**
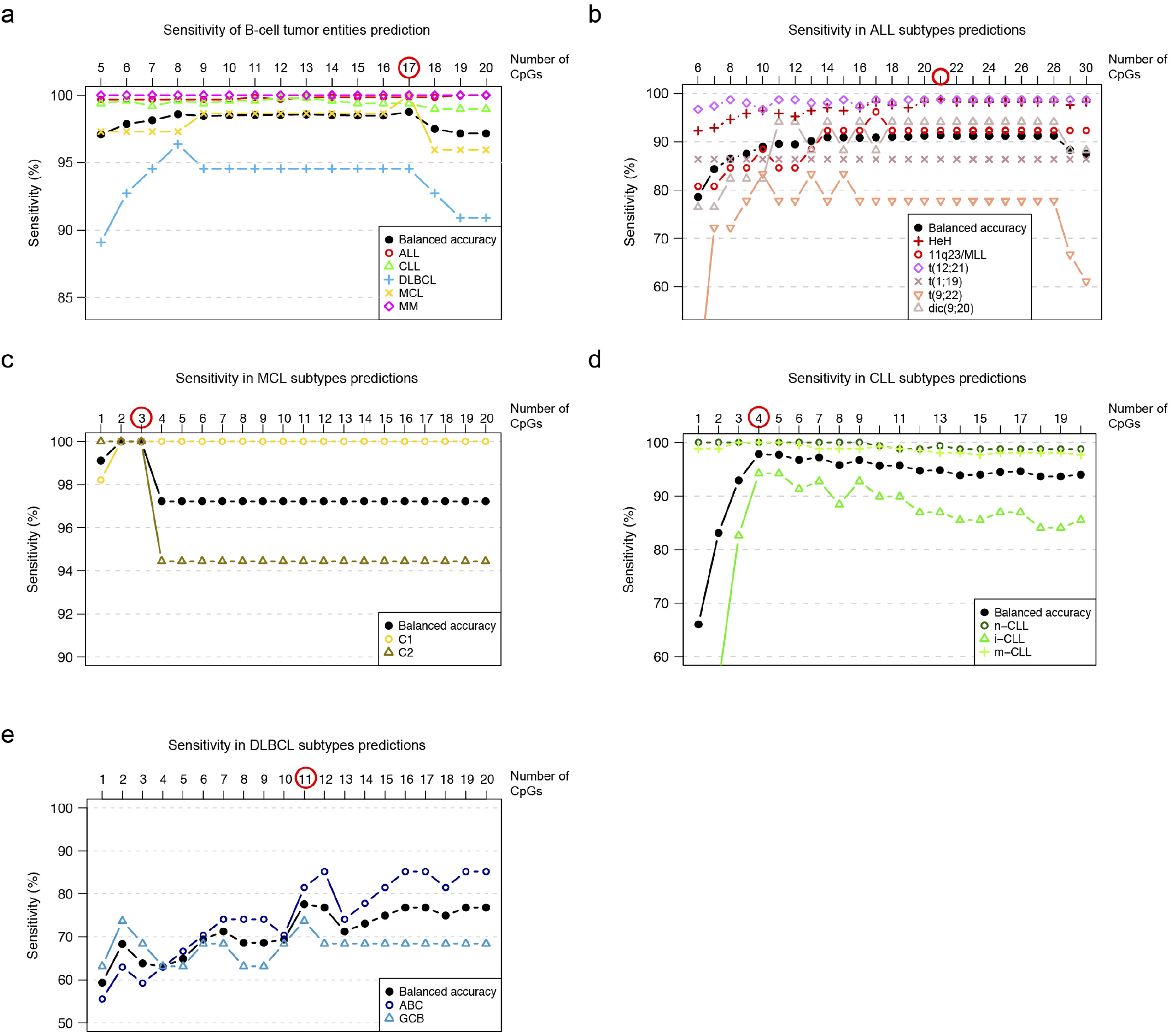
**a,** Sensitivity of the pan-B-cell diagnostic algorithm for the classification of an unknown B-cell tumor into ALL, MCL, CLL, DLBCL or MM while incrementing the number of CpGs used in the classifier algorithm (first step of Fig. 3A). The number of CpGs selected for the predictor is selected by maximizing the highest balanced accuracy and is indicated with a red circle. This strategy was applied also in the remaining 4 predictors to classify B-cell tumor subtypes in panels b, c, d, and e, (second step of Fig. 3A). Each B-cell tumor is represented with different shapes and colors. ALL, acute lymphoblastic leukemia; MCL, mantle cell lymphoma; CLL, chronic lymphocytic leukemia; DLBCL, Diffuse large B cell lymphoma; MM, multiple myeloma **b,** Sensitivity of predictor 2 for the pan-B-cell diagnostic algorithm (predictor 2 of Fig. 3a) for the classification of ALL into the subtypes HeH, 11q23/MLL, t(12;21), t(1;19), t(9;22) and dic(9;20) while incrementing the number of CpGs. **c,** Sensitivity of predictor 3 of the pan-B-cell diagnostic algorithm (predictor 3 of Fig. 3a) for the classification of MCL into the subtypes C1 or C2 while incrementing the number of CpGs. **d,** Sensitivity of predictor 4 of the pan-B-cell diagnostic algorithm (predictor 4 of Fig. 3a) for the classification of CLL into the subtypes n-CLL, i-CLL or m-CLL while incrementing the number of CpGs. **e,** Sensitivity of predictor 5 of the pan-B-cell diagnostic algorithm (predictor 5 of Fig. 3a) for the classification of DLBCL into the subtypes ABC and GCB while incrementing the number of CpGs.

**Extended Data Fig.4.**
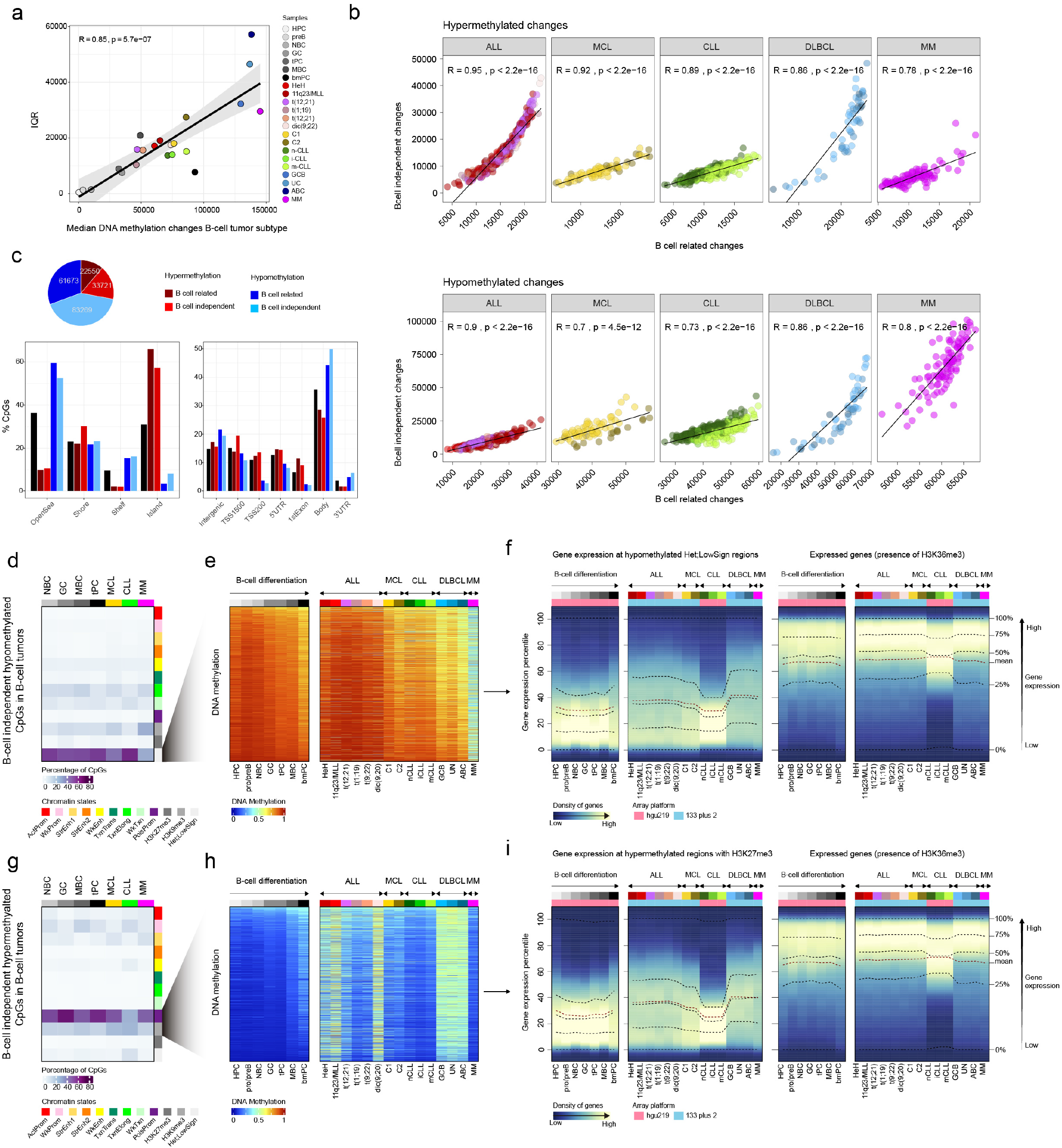
**a,** Variability of DNA methylation changes (IQR) in normal neoplastic B cells against the median number of DNA methylation changes. **b,** Correlations of B-cell independent DNA methylation changes in B-cell tumors against B-cell related changes for hypermethylation (top) and hypomethylation (bottom). **c,** Number of B-cell related or B-cell independent hyper- or hypomethylation in B-cell tumors (Methods). **d,** B-cell independent CpGs losing DNA methylation in B-cell tumors and the percentages in each chromatin state in normal and neoplastic B-cells. **e,** Example CpGs from **d,** in normal and neoplastic B-cells. **f,** Density of genes distributed along gene expression percentiles of genes associated with B-cell independent CpGs losing DNA methylation in B-cell tumors in each B-cell tumor subtype. Expressed genes are displayed at right as control (with presence of H3K36me3). Means within each B-cell subpopulation as well as B-cell tumors are represented. **g,** B-cell independent CpGs gaining DNA methylation in B-cell tumors and the percentages in each chromatin state in normal and neoplastic B-cells. **h,** Example CpGs from g, in normal and neoplastic B-cells. **i,** Density of genes distributed along gene expression percentiles of genes associated with B-cell independent CpGs gaining DNA methylation in B-cell tumors in each B-cell tumor subtype. Expressed genes are displayed at right as control (with presence of H3K36me3). Means within each B-cell subpopulation as well as B-cell tumors are represented.

**Extended Data Fig.5.**
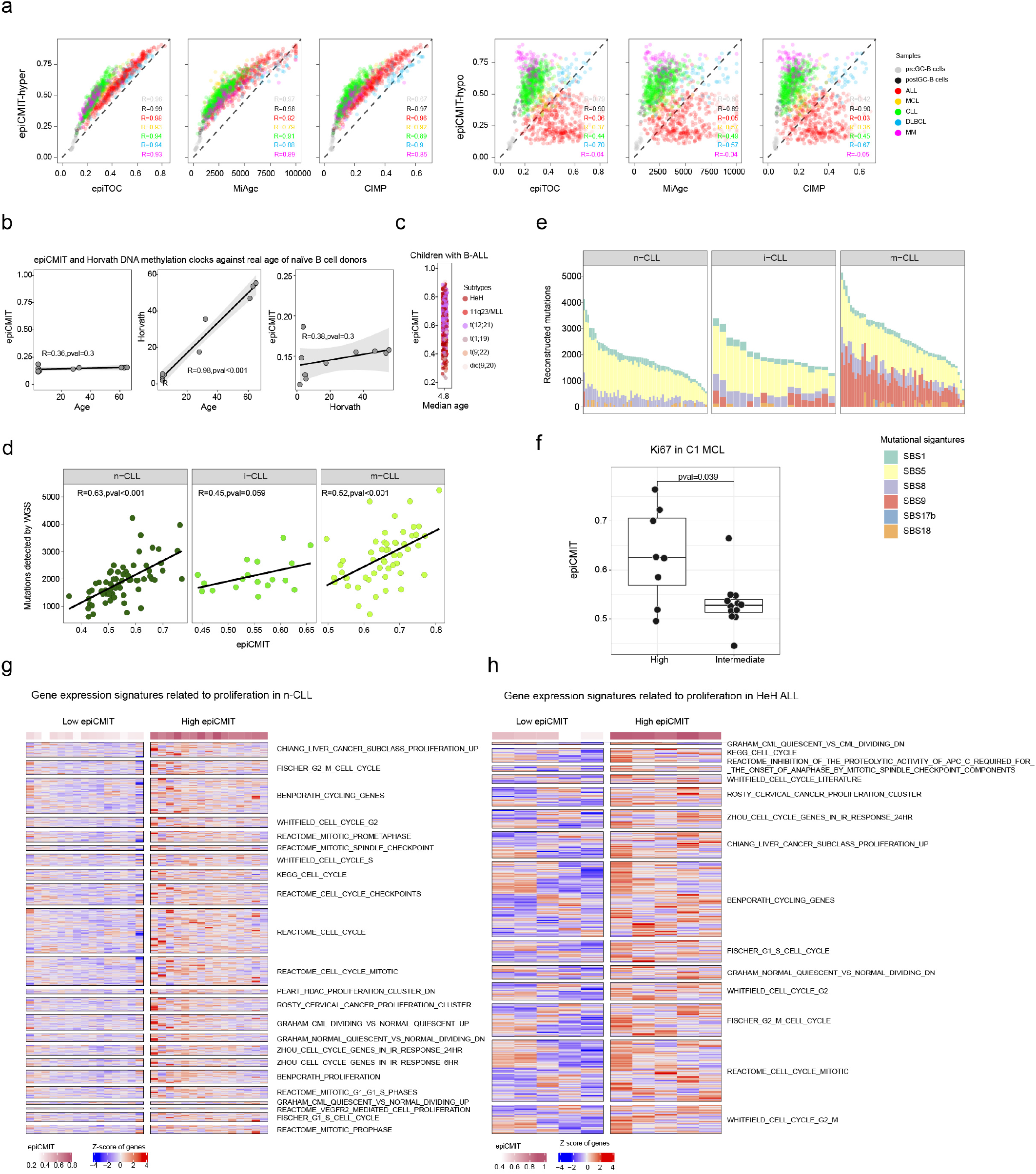
**a,** Correlations between epiCMIT-hyper and epiCMIT-hypo with other hypermethylation-based mitotic clocks including epiTOC (epigenetic Timer for Cancer risk) and MiAge as well as the CIMP (CpG hypermethylator phenotype, not formally conceived as mitotic clock). epiCMIT, epigenetically-determined Cumulative MIToses. **b,** Poor effect of age in epiCMIT in normal B cells. epiCMIT correlation with age in naïve B cells from healthy infants, young-adults and older adults. Horvath model correctly predicted the real age. epiCMIT and Horvath are poorly correlated. **c,** Poor effect of age in epiCMIT in B-ALL. Wide epiCMIT spectrum in children with BALL. **d,** Correlation between the number of mutations detected by WGS against epiCMIT in CLL subtypes with different cellular origin (n-CLL, i-CLL and m-CLL). **e,** Mutational signatures for 138 CLL samples with available WGS and DNA methylation data divided into different subtypes of CLL with different cellular origin, namely n-CLL, i-CLL and m-CLL. **f,** Correlation of epiCMIT with Ki67 in nodal MCL patients. **g,** Gene expression signatures resulting after GSEA analysis related to proliferation in CLL samples with low and high epiCMIT. **h,** Gene expression signatures resulting after GSEA analysis related to proliferation in ALL samples with low and high epiCMIT.

**Extended Data Fig.6.**
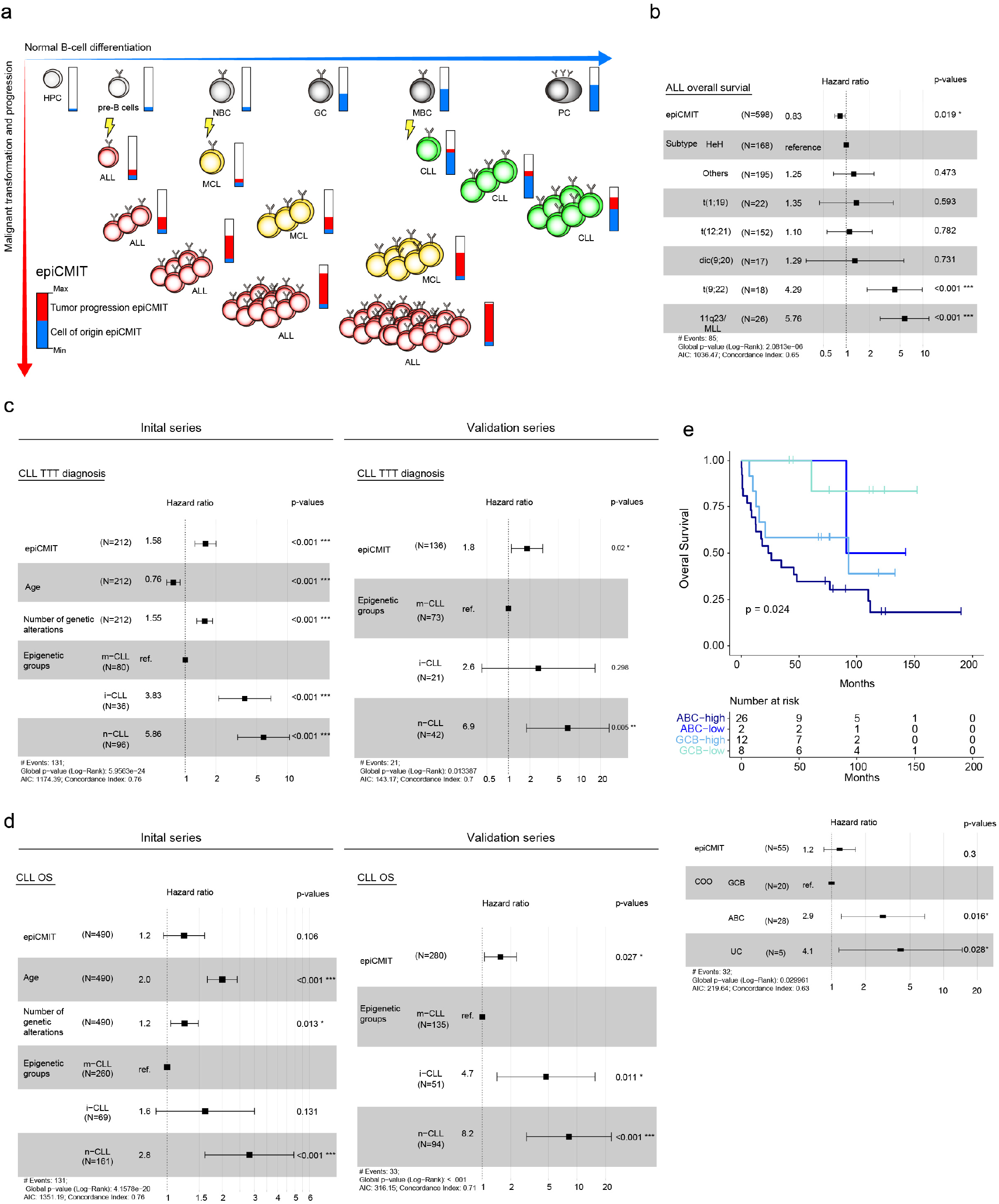
**a,** epiCMIT interpretation in normal and neoplastic B cells. Mitotic cell division occurs in normal B cell development and DNA methylation changes accumulate at repressed regions (blue component of the epiCMIT on the x axis). B-cell tumors arise from different maturation stages, and thus they contain different baseline epiCMIT from the cell of origin from which they originate (blue bar in B-cell tumors). When B-cell tumors progress, they acquire additional DNA methylation changes at repressed regions (shown as red component of the epiCMIT). Notably, these new DNA methylation changes occur in both axes, i.e. changes occurring in normal B cell maturation as well as new B-cell independent changes. This concept is shown with real data at Fig. 4b. **b,** Multivariate cox regression model for overall survival in ALL for epiCMIT against cytogenetic groups. Hazard ratio for epiCMIT correspond to 0.1 increments. **c,** Multivariate cox regression model for time to first treatment in CLL samples near diagnosis (30 months) for epiCMIT against age, number of driver alterations and epigenetic groups based on different cellular origin. At right is shown the multivariate cox regression model for validation series. Hazard ratio for epiCMIT correspond to 0.1 increments. **d,** Multivariate cox regression model in CLL for overall survival for epiCMIT against age, number of driver alterations and epigenetic groups for the initial and validation series (left and right, respectively). Treated and untreated patients were included in the validation series. Hazard ratio for epiCMIT correspond to 0.1 increments. **e,** Kaplan-Meyer curves for DLBCL separated by ABC and GCB groups and low and high epiCMIT groups based on maxstat rank statistics. Multivariate cox regression model treating epiCMIT as continuous variable is shown at bottom. Hazard ratio for epiCMIT correspond to 0.1 increments.

**Extended Data Fig.7.**
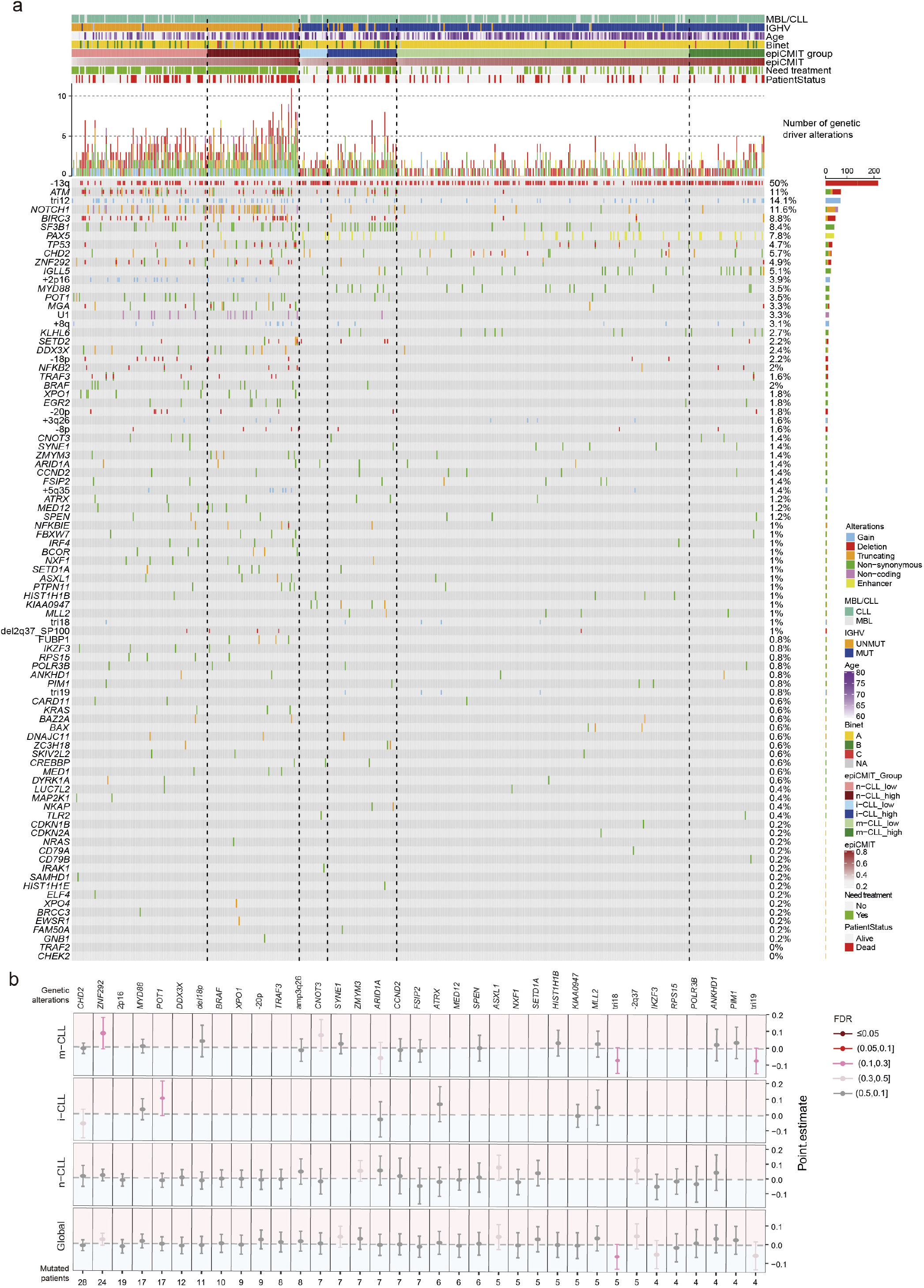
**a,** Oncoprint showing all genetic driver alterations considered in CLL grouped by epigenetic subgroups and ordered according to increasing levels of epiCMIT (from left to right within each epigenetic subgroup). Other clinicobiological features including MBL or CLL, IGHV status, Age, Binet stage, epiCMIT subgroups based on Fig. 5C, need for treatment and patient status are shown. Distinct genetic driver alterations are depicted with different colors and shapes. The percentage of mutated patients as well as barplots showing the number of mutated patients for each alteration is shown at right. **b,** Driver genetic alterations without clear associations with epiCMIT. Analyses were done globally for all CLL samples (although adjusted by epigenetic groups) as well as within each epigenetic subgroup. Positive point estimates relate to positive associations with epiCMIT. 95% confidence intervals are shown with colors coding for FDR corrections. Number of patients with each genetic driver alteration is shown at bottom.

